# A key role of pectin demethylation-mediated cell wall Na^+^ retention in regulating differential salt stress resistance in allotetraploid rapeseed genotypes

**DOI:** 10.1101/2023.09.09.556983

**Authors:** Ting Zhou, Peng-jia Wu, Jun-fan Chen, Xiao-qian Du, Ying-na Feng, Cai-peng Yue, Jin-yong Huang, Ying-peng Hua

**Affiliations:** School of Agricultural Sciences, Zhengzhou University, Zhengzhou, 450001, China; Zhengzhou Key Laboratory of Quality Improvement and Efficient Nutrient Use for Main Economic Crops, Zhengzhou 450001, China; School of Life Sciences, Zhengzhou University, Zhengzhou, 450001, China

**Author notes:** To whom correspondence should be addressed: Ying-peng Hua. Email address for each author: Ting Zhou Peng-jia Wu Jun-fan Chen Xiao-qian Du Ying-na Feng Cai-peng Yue Jin-yong Huang Ying-peng Hua.

**Keywords:** Allotetraploid rapeseed, cell wall, multiomics, pectin methylation, pectin methyl esterase, salt stress resistance

## Abstract

Allotetraploid rapeseed (*Brassica napus* L.) is highly susceptible to salt stress, a worldwide limiting factor that causes severe losses in seed yield. Genetic variations in the resistance against salt stress found in rapeseed genotypes emphasizes the complex response architecture. Westar is ubiquitously used as a major transgenic receptor, and ZS11 is widely grown as a high production and good quality cultivar. In this study, Westar was identified to outperform than ZS11 under salt stress. Through cell component isolation, non-invasive micro-test, X-ray energy spectrum analysis, and ionomic profiling characterization, pectin demethylation was found to be a major regulator for differential salt resistance between Westar and ZS11. Integrated analyses of genome-wide DNA variations, differentially expression profiling, and gene co-expression network identified *BnaC9.PME47*, encoding pectin methyl esterase, as a positive regulator mainly responsible for salt stress resistance. *BnaC9.PME47*, located in two reported QTLs regions for salt resistance, was strongly induced by salt stress and localized on the cell wall. Natural variation of the promoter regions conferred higher expression of *BnaC9.PME47* in Westar than in other salt-sensitive rapeseed genotypes. Loss-of-function of *AtPME47* resulted in the hypersensitivity of Arabidopsis plants to salt stress. This study facilitates a more comprehensive understanding of the differential morpho-physiological and molecular responses to salt stress and abundant genetic diversity in rapeseed genotypes, and the integrated multiomics analyses provide novel insights regarding the rapid dissection of quantitative trait genes responsible for nutrient stresses in plant species with complex genomes.

## Introduction

Sodium ion (Na^+^) is one of the most abundant chemical elements in the earth’s crust **(Nieves-Cordones et al., 2016)**. Particularly, in arid and semi-arid arable regions, excessive Na^+^ easily result in severe soil salinization, a severe abiotic stress factor that hinders crop yield and quality **(Setia et al., 2013)**. Soil salinity usually leads to adverse effects, including imbalanced essential nutrient homeostasis, damaged membrane structures, and dysfunctional antioxidant systems **(Munns and Gilliham, 2015)**. Exploiting saline soils to grow glycophyte crops, increasing yield production, is an important route for meeting the ever-increasing demands for foods across the globe. Therefore, understanding the mechanisms underlying salt stress resistance is a crucial prerequisite to develop agriculture crops with improved salinity resistance **(Negrão et al., 2017; Shao et al., 2020)**.

In general, plants have evolved multiple strategies to counter salinity stress through regulating Na^+^ homeostasis: (i) reduce root Na^+^ absorption, (ii) increase intracellular Na^+^ compartmentation and extrusion, and (iii) enhance xylem Na^+^ unloading **(Assaha et al., 2017)**. Recently, more and more studies reveal that plants respond to salt stress and cope with the resulting damages by altering the biosynthesis and deposition of the main cell wall components to prevent water loss and decrease the entry of surplus ions into the plants. Salt stress may cause cell wall softening, notably through its effect on pectins **(Feng et al., 2018)**. Excessive Na^+^ in apoplastic regions competes with calcium (Ca^2+^) to bind to pectins, which in turn disturbs pectin cross-linking **(Byrt et al., 2018)**.

Pectins are typically secreted in a highly methyl-esterified form and then selectively de-esterified by pectin methylesterases (PMEs). PMEs (EC 3.1.1.11) function in de-esterification of the methylated carboxyl group (COOCH_3_) of pectin to form elastic pectins and accompany MeOH generation. Pectins that are de-methylesterified can be cross-linked by divalent cations, such as Ca^2+^, to form egg-box structures and thus promote cell wall stiffness **(Hocq et al., 2017)**. Currently, *AtPME31* was found to positively regulate salt stress tolerance in Arabidopsis **(Yan et al., 2018)**. Mutation of *OsTSD2*, which encodes a pectin methyltransferase, leads to reduced salt tolerance in rice **(Fang et al., 2019)**. However, few PMEs were functionally identified to be involved in the responses of ployploid crops to salt stress.

Rapeseed (*Brassica napus* L., A_n_A_n_C_n_C_n_, 2n = 4x= 38) is an allotetraploid crop species originating from the diploid ancestors *B. rapa* and *B. oleracea*, and is the third most important oilseed crop after palm and soybean **(Song et al., 2020)**. Salt stress severely hinders the growth, development, seed yield production of rapeseed **(Shah et al., 2021)**. Genetic variations in the resistance against salt stress found in rapeseed genotypes emphasizes the complex response architecture **(Ibrahim et al., 2022; Wan et al., 2022)**. The breeding of salt-resistant rapeseed germplasm is a cost-effective and environmentally friendly strategy for the efficient utilization of saline soils. However, few genes that can be used for the genetic improvement of salt stress resistance have been isolated and functionally validated in allotetraploid rapeseed.

Zhongshuang 11 (ZS11) and Westar, both of whose genome information have been released, are two rapeseed cultivars widely used in fundamental studies and practical production of rapeseed **(Song et al., 2020)**. According to previous studies, Westar is identified as a tolerant cultivar under salt stress **(Yong et al., 2015)**, while ZS11 is highly sensitive to salinity **(Ibrahim et al., 2022; Wan et al., 2022)**. However, whether Westar had different responses to salt stress from ZS11 was not reported before; moreover, if different, the corresponding physiological, genetic, and molecular mechanisms remain to be elucidated.

In this study, first, we found that Westar was more tolerant against salt stress than ZS11. Then, we combined the ionomic analysis, genomic polymorphism characterization, differential transcriptional and phytohormone profiling, and functional validation to reveal the physiological, genomic, and molecular bases of differential salt stress resistance between Westar and ZS11. This study provides theoretical reference about the selection of transgenic receptors for studying rapeseed salt resistance, and also provide new gene resources for the genetic improvement of salt stress resistance in ZS11. The integrated multiomics analyses provide novel insights regarding the rapid dissection of quantitative trait genes responsible for nutrient stresses in plant species with complex genomes.

## Materials and Methods

### Plant materials and growth conditions

ZS11, developed by Oil Crops Research Institute of Chinese Academy of Agricultural Sciences (Wuhan, China), is a wide-grown semi-winter rapeseed variety with high oil content and seed production, and low erucic acid and glucosinolate content **(Sun et al., 2017)**. ZS11 shows strong resistance against diverse nutrient stresses, such as boron deficiency **(Hua et al., 2016)** and cadmium toxicity **(Zhang et al., 2019)**. Westar, developed by the Canadian Saskatoon Agricultural Research Station (Saskatoon, Canada), is a spring rapeseed cultivar that is widely used as a transgenic receptor **(Cardoza and Stewart, 2003)**, while it shows highly sensitivity to multiple abiotic and biotic stresses, such as boron deficiency **(Hua et al., 2016)** and blackleg disease **(Andreasson et al., 2001)**.

The rapeseed plants were hydroponically grown in an illuminated growth chamber using the Hoagland and Arnon nutrient solution. The setup of rapeseed growth conditions was as follows: light intensity of 150 μmol m^-2^ s^-1^, temperature of 24°C daytime/22°C night, light period of 16-h photoperiod/8-h dark, and relative humidity of 50% **(Zhou et al., 2021)**.

*Arabidopsis thaliana* L. Heynh. Ecotype Columbia (Col-0), *pme47-1* (SALK_094562) and *pme47-2* (SALK_104093) were obtained from the Arabidopsis Biological Resource Center (ABRC). Homogeneity of *pme47-1* and *pme47-1* were identified by T-DNA insertion-bases PCR using the specific primers as indicated in **Supplementary Table S1**. Seeds were surface sterilized and sown on 1/2 MS medium plates (Murashige and Skoog salts including vitamins and 1% (w/v) sucrose). For salt treatment, uniform 10-d-old Arabidopsis seedlings were transferred to 1/2 MS medium plates with 100 mM NaCl for 5 d until sampling.

### Analysis of plant morphology and cell ultrastructure

The roots of rapeseed/Arabidopsis seedlings were subjected to an image scanner for the analysis of total root length, root volume, root surface area, root average diameter, and root tip number using WinRHIZO Pro (Regent Instruments, QC, Canada).

Intracellular ultrastructure of leaf pieces (approximately 1 mm^2^) from rapeseed plants, which were treated with 200 mM NaCl for 5 d, were determined by transmission electron microscopy (H-7650; Hitachi, Tokyo, Japan). A SEM (JSM-6390/LV, JEOL, Tokyo, Japan) was used to characterize the stomatal structure. The sample pieces were prepared as described previously **(Dittmer et al., 2021)**, and each test had at least five independent biological replicates.

### Determination of physiological parameters and quantification of mineral elements

SPAD (Soil and Plant Analyzer Development) estimation of leaves from rapeseed plants, which were treated with 200 mM NaCl for 5 d, were determined using a SPAD-502 chlorophyll meter (Konica Minolta, Tokyo, Japan). The concentrations of malondialdehyde (MDA), which was extracted using thiobarbituric acid, were assayed spectrophotometrically at the wavelengths of 450 nm, 532 nm, and 600 nm **(Kotula et al., 2020)**. Proline concentrations were determined spectrophotometrically at 520 nm using the ninhydrin method **(Deng et al., 2016)**. To detect hydrogen peroxide (H_2_O_2_), 3,3’-Diaminobenzidine (DAB) staining was used as previously described **(Ouyang et al., 2010)**. The superoxide anion (O_2_ ^-^) was detected with the NBT staining method **(Kaur et al., 2016)**.

Sampled rapeseed plants, which were treated with 200 mM NaCl for 5 d, were divided into roots and shoots and were oven-dried at 65°C until a constant weight was achieved. The dried tissues were subsequently transferred to a HNO_3_/HClO_4_ mixture (4:1, v/v) at 200°C until they were completely digested **(Zhang et al., 2019)**. The samples were then diluted with deionized water, and the concentrations of metal elements were quantified by inductively coupled plasma mass spectrometry (ICP-MS; NexION^TM^ 350X, PerkinElmer).

### TEM coupled with X-ray spectrum analysis of subcellular Na^+^ distribution

To examine the relative differences in vacuolar Na^+^ compartmentation between Westar and ZS11, a transmission electron microscope (H800, Hitachi, Tokyo, Japan) coupled with an EDAX-910 energy-disperse X-ray analyzer was used to determine relative abundances of shoot vacuolar Na^+^. The rapeseed samples were prepared following our previous study **(Hua et al., 2023)**. The accelerating voltage was set as 120 kV with a take-off angle of 25°C, and the counting time was set to 60 s. The relative Na^+^ abundances are expressed as counts per second (CPS) of the Na^+^ peak following subtracting the background values **(Lv et al., 2012)**.

### Identification and characterization of genomic DNA variations

Functional annotation of the genome-wide DNA variants between Westar and ZS11, whose genome sequences were download from the BnIR database (https://yanglab.hzau.edu.cn/BnIR), was performed with ANNOVAR, and for gene and region annotations, UCSC was used. Gene ontology (GO) and Kyoto Encyclopedia of Genes and Genomes (KEGG) analyses were performed using Panther Classification System (http://www.pantherdb.org/) **(Mi et al., 2013)** and KEGG Mapper (https://www.kegg.jp/kegg/pathway.html) **(Kanehisa and Sato, 2020)** to annotate the biological function and pathways of target genes. The coding sequences and promoter regions of target genes were amplified using the high-fidelity Phusion NEB DNA polymerase. The primers are listed in **Supplementary Table S1**.

### RNA extraction and transcriptome sequencing

Total RNA from the above fresh rapeseed shoots, including 12 samples, was extracted using the Invitrogen TRIzol^®^ Reagent (Invitrogen, CA, USA), and genomic DNA was removed by TaKara DNase I. RNA quality and quantification was determined by 2100 Bioanalyser (Agilent, CA, USA) using NanoDrop 2000 (Thermo Fisher Scientific, Massachusetts, USA), respectively. Only high-quality RNA samples (OD_260_/OD_280_=1.8∼2.2, OD_260_/OD_230_ ≥ 2.0, RIN ≥ 8.0, 28S:18S ≥ 1.0) were used to construct sequencing libraries.

Transcriptome libraries were prepared following the instructions of TruSeq^TM^ RNA sample preparation Kit (Illumina, San Diego, CA, USA). Briefly, mRNA was isolated based on poly-A selection using the oligo(dT) beads, and fragmentation was performed in fragmentation buffer. Then, the synthesis of double-stranded cDNA was performed using a SuperScript double-stranded cDNA synthesis kit (Invitrogen, CA, USA) with random hexamer primers (Illumina). Next, the synthesized cDNA was subjected to end-repair, phosphorylation, and ‘A’ base addition according to Illumina’s library construction protocol. Libraries were size selected (200-300 bp) using 2% Low Range Ultra Agarose. After quantification by TBS380 fluorometer, the paired-end (2 × 150 bp read length) RNA-seq sequencing library was sequenced by the Illumina HiSeq Xten system. The raw paired-end reads were trimmed and quality controlled by SeqPrep (https://github.com/jstjohn/SeqPrep) and Sickle (https://github.com/najoshi/sickle) programs with default parameters. Next, clean reads were separately aligned to reference genome using TopHat v2.0.0 (http://tophat.cbcb.umd.edu/) software.

Expression levels of transcripts were calculated as fragments per kilobase of exon per million mapped reads (FRKM) values via RSEM (http://deweylab.biostat.wisc.edu/rsem/). Fold-change ≥ 2 and false discovery rate (FDR) < 0.05 were used to identify differentially expressed genes (DEGs). R statistical package, EdgeR (Empirical analysis of Digital Gene Expression (http://www.bioconductor.org/packages/2.12/bioc/html/edgeR.html), was used for differential expression analysis. GO and KEGG enrichment analysis of DEGs was performed by GOATOOLS (https://github.com/tanghaibao/Goatools) and KOBAS (http://kobas.cbi.pku.edu.cn/home.do) at Bonferroni-corrected *P*-value ≤ 0.05 compared with the background whole-transcriptome.

### Molecular characterization of candidate genes

The genes in *Brassica* species are named following the criterion of genus (one capital letter) + plant species (two lowercase letters) + chromosome (followed by a period) + name of corresponding homologs in *A. thaliana* **(Zhou et al., 2020)**. The target genes were physically mapped onto rapeseed chromosomes using MapGene2Chromosome v2.1. KaKs_calculator was used to calculate the nonsynonymous (Ka) and synonymous (Ks), substitution rates, and Ka/Ks based on the pairwise CDS and amino acid alignment with the yn00 method **(Yang and Nielsen, 2000)**. According to the Darwin’s evolution theory, it is proposed that Ka/Ks > 1.0 means positive selection, while Ka/Ks < 1.0 indicates purifying selection, and Ka/Ks = 1.0 denotes neutral selection. Tissue-specific expression profiles of target genes were retrieved from the *Brassica napus* information resource (BnIR) database (https://yanglab.hzau.edu.cn/BnIR). To identify putative *cis*-acting regulatory elements, two-kb of upstream genomic sequences from the start codon (ATG) of target genes were analyzed by PLACE v. 30.0 (http://www.dna.affrc.go.jp/PLACE/) **(Higo et al., 1999)** and plantCARE (http://bioinformatics.psb.ugent.be/webtools/plantcare/html/) **(Lescot et al., 2002)**.

The *pCAMBIA1300*-GFP expression vector fused with target genes were introduced into tobacco leaf using *Agrobacterium tumefaciens* strain GV3101, and the plasma membrane of tobacco leaf cells were stained by FM4-64. For the plasmolysis of tobacco leaf cells, the leaves were treated by 5% NaCl for 5 min **(Chen et al., 2017)**. Fluorescence was observed using a Nikon C2-ER confocal laser scanning microscope (Nikon, Japan) following emission filter sets at 510 nm (GFP) and 580 nm (RFP), and excitation was achieved at 488 nm (GFP) and 561 nm (RFP).

### Reverse transcription-quantitative polymerase chain reaction (RT-qPCR) assays

RT-qPCR assays were performed to examine the relative expression of target genes. After removing genomic DNA from RNA samples with RNase-free DNase I, total RNA was used as RT templates for cDNA synthesis using the TaKaRa PrimeScript^TM^ RT Reagent Kit Eraser. The RT-qPCR assays were conducted using TaKaRa SYBR^®^*Premix Ex Taq*^TM^ II under a Bio-Rad’s C1000 touch Thermal Cycler of CFX96^TM^ Real-time PCR detection System.

The RT-qPCR program was as follows: 95°C for 3 min, 40 cycles of 95°C for 10 s, and 60°C for 30 s. To analyze the primer gene-specificity, the melt curve was plotted as follows: 95°C for 15 s, 60°C for 1 min, and 60-95°C for 15 s (+0.3°C/cycle). The gene expression levels were calculated with the 2^-ΔΔC^*_T_* method **(Livak and Schmittgen, 2001)** using *BnaEF1-*α **(Maillard et al., 2016)** and *BnaGDI1* **(Yang et al., 2014)** as the internal reference. The primers used in this study are listed in **Supplementary Table S1**.

### Measurement of K^+^ and Na^+^ fluxes by non-invasive micro-test technology

The net Na^+^ and K^+^ fluxes were measured using non-invasive micro-test technology (Younger USA LLC, Amherst, MA, USA) **(Feng et al., 2015)**. In order to measure net Na^+^ and K^+^ fluxes, rapeseed seedlings were treated with 200 mM NaCl solution (pH 8.0) for 24 h, then Na^+^ and K^+^ effluxes were measured at the meristem zone of primary roots (∼500 μm from the root tip). The detailed experimental procedures were performed according to a previous study **(Liu et al., 2019)**. Data were collected at 6-s intervals during the measurements, and each point represents the average of eight biological samples **(Yang et al., 2019)**.

### Fractionation of cell wall components and analysis of Fourier-transform infrared spectrometry

Extraction of cell walls and isolation of pectin was performed in accordance with the method described by **Zhu et al. (2015)**. Briefly, the rapeseed shoots were ground to fine power in liquid nitrogen. Crude cell walls were extracted using ice-cold 75% alcohol, followed by acetone, methanol:chloroform (1:1, v/v), and methanol, and were then lyophilized and stored at 4 °C until use. The supernatant, which was collected via hot-water extraction, was collected as pectins.

The relative abundances of functional groups in the lyophilized cell walls, including carboxyl (COO^−^) groups, were analysed using Fourier-transform infrared spectrometry (FTIR; Vertex 70, Bruker Optics, Ettlingen, Germany) as described by **Zhou et al. (2017)**. For each sample, five biological replicates were examined under the same conditions.

### Determination of phytohormone profiling

Seven phytohormone classes (indole-3-acetic acid, abscisic acid, salicylic acid, gibberellin, cytokinin, brassinosteroid, and jasmonates) were tested using a HPLC-MS/MS method **(Almeida Trapp et al., 2019)**. Tubes containing 100 mg of fine powder of rapeseed shoots and roots were kept at −80^◦^C, and transferred to liquid nitrogen before the extraction. Each sample was mixed with 750 μL prechilled extraction buffer (methanol:water:acetic acid, 80:19:1, v/v/v) supplemented with internal standards to obtain the phytohormones extract **(Almeida Trapp et al., 2014)**. The phytohormone concentrations were determined by ultra-fast liquid chromatography-electrospray ionization tandem mass spectrometry (UFLC-ESI-MS).

### Data availability

All data supporting the findings of this study are available within the paper and within its supplementary data published online. The raw data of the high-throughput transcriptome sequencing have been deposited in the NCBI Sequence Read Archive (SRA) under the Bioproject accession no. PRJNA952357.

## Results

### Differential growth performance of the shoots and roots between Westar and ZS11 under salt stress

Prior to the identification of differential salt stress resistance between Westar and ZS11, we first tested their seed germination rates under control and salt stress conditions **(Fig. 1A-C)**. It is obvious that salt stress significantly repressed the seed germination in both Westar and ZS11 compared with the salt-free condition **(Fig. 1A)**. In terms of the differences in the seed germination between the two genotypes, Westar had a higher germination rate than ZS11 **(Fig. 1B)**, whose seminal root lengths were also smaller than those of Westar **(Fig. 1C)**. Under control condition, the two rapeseed genotypes showed no marked differences in growth performance; however, at 200 mM NaCl for 5 d, the cotyledons of ZS11 showed severe chlorosis than Westar **(Fig. 1D)**, particularly in the cotyledons and 1st/2nd euphylls **(Fig. 1E)**. We also observed that salt stress had a more pronounced effect on stomatal size and morphology in ZS11 than in Westar **(Fig. 1F, G)**. Determination of leaf SPAD values showed that ZS11 suffered from more severe leaf chlorosis than Westar under salt stress **(Fig. 1H)**, which also induced a higher proportion of water loss in the shoots of ZS11 than in the shoots of Westar **(Fig. 1I)**. Under salt stress, the concentrations of MDA and proline in both Westar and ZS11 were significantly increased than under control **(Fig. 1J)**. The proline concentrations in Westar were obviously higher than in ZS11, which had higher MDA concentrations **(Fig. 1K)**. Quantitative determination of reactive oxygen species showed that ZS11 had higher levels of H_2_O_2_ and O_2_^-^ than Westar **(Fig. 1L, M)**; moreover, the DAB and NBT staining also confirmed that ZS11 accumulated more reactive oxygen species than Westar **(Fig. 1N O)**.

**Fig. 1.**
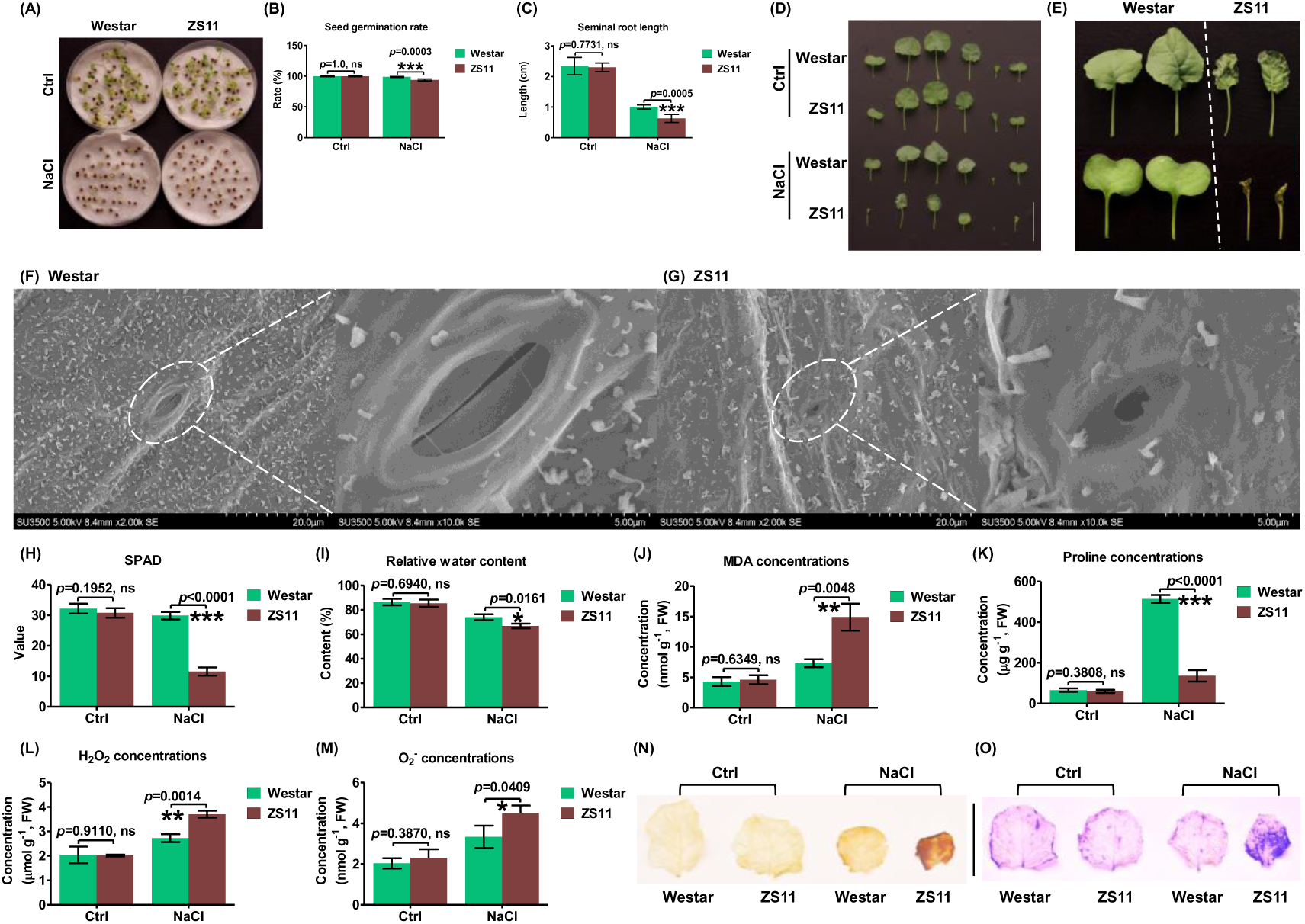
Different shoot growth performance between the salt-resistant Westar and the salt-sensitive ZS11 under control and salt stress conditions. **(A)** Growth performance of germinated seeds; **(B-C)** seed germination rates and hypocotyl lengths of germinated seedlings. Plump rapeseed seeds of Westar and ZS11 were sowed hydroponically under salt stress condition (200 mM NaCl) for 5 d. **(D)** Overview of the shoot growth performance under NaCl-free and salt stress conditions. Scale bar = 5 cm. **(E)** Growth performance of the cotyledons and 1^st^/2^nd^ euphylls under salt stress. Scale bar = 2 cm. **(F-G)** Stomata morphology of Westar **(F)** and ZS11 **(G)** under salt stress. Scale bar = 5 μm. **(H-O)** Leaf SPAD values **(H)**, relative water content **(I)**, malondialdehyde (MDA) concentrations **(J)**, proline concentrations **(K)**, H2O2 concentrations **(L)**, O2^-^ concentrations **(M)**, and DAB **(N)** and NBT **(O)** staining in the shoots of Westar and ZS11 under NaCl-free and salt stress conditions. Scale bar = 5 cm. Ctrl, control. Data are presented as the mean (n=5) ± s.d. Significant differences (*, *p*<0.05; **, *p*<0.01; ***, *p*<0.001) were determined by unpaired two-tailed Student’s *t*-tests between two groups using the SPSS 17.0 toolkit. ns, not significant.

Further, we compared the differential responses of rapeseed roots to salt stress between Westar and ZS11 through investigating their root architecture system **(Fig. 2)**. Similar to our previous study **(Zhang et al., 2019)**, the root growth performance of Westar and ZS11 did not show obvious differences under control condition **(Fig. 2A)**. However, consistent with the better shoot growth performance in Westar under salt stress, the roots of Westar also grew much stronger than those of ZS11 **(Fig. 2B)**. In detail, when exposed to salt stress, relative to ZS11, Westar presented larger primary root lengths **(Fig. 2C)**, total root lengths **(Fig. 2D)**, root surface areas **(Fig. 2E)**, and root volumes **(Fig. 2F)**, while these two rapeseed genotypes did not show remarkable differences in the average root diameters **(Fig. 2G)** and root tip number **(Fig. 2H)**.

**Fig. 2.**
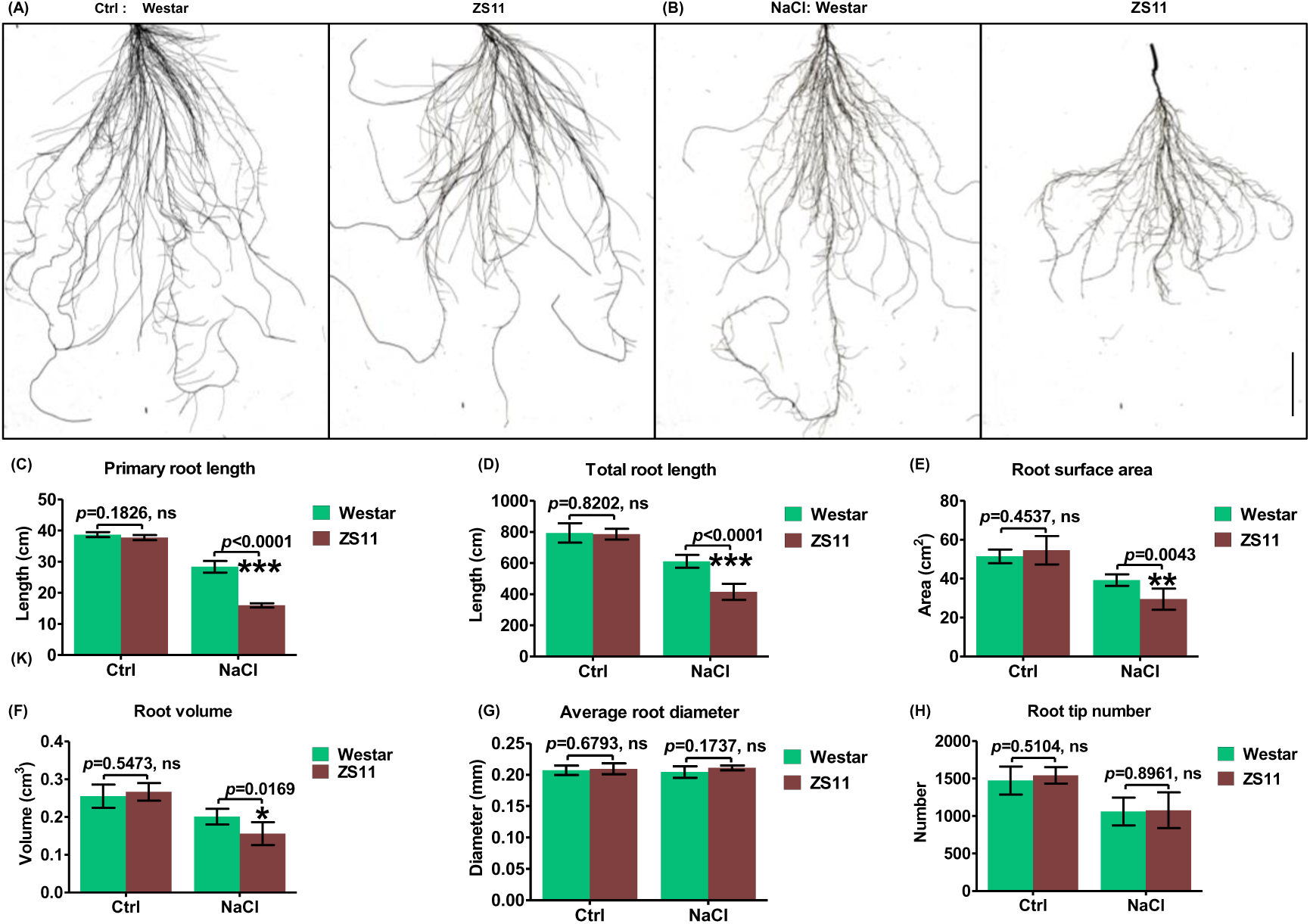
Different root growth performance between the salt-resistant Westar and the salt-sensitive ZS11 under control and salt stress conditions. **(A-B)** Growth performance of the roots between Westar and ZS11 under control **(A)** and salt stress **(B)** conditions. **(C-H)** Primary root length **(C)**, total root length **(D)**, root surface area **(E)**, root average diameter **(F)**, root volume **(G)**, and root tip number **(H)** of Westar and ZS11 under control and salt stress conditions. Scale bar = 5 cm. Uniform rapeseed plants after seed germination were grown hydroponically under NaCl-free conditions for 10 d, and then transferred to 200 mM NaCl for 5 d. Ctrl, control. Data are presented as the mean (n=5) ± s.d. Significant differences (*, *p*<0.05; **, *p*<0.01; ***, *p*<0.001) were determined by unpaired two-tailed Student’s *t*-tests between two groups using the SPSS 17.0 toolkit. ns, not significant.

### Pectin methylation-mediated cell wall Na^+^ retention might be involved in the differential salt stress resistance between Westar and ZS11

Based on the significant morpho-physiological differences between the salt-resistant genotype Westar and the salt-sensitive genotype ZS11, ICP-MS was employed to reveal the ionomic basis of differential salt stress resistance. Firstly, total Na^+^ concentrations of whole plants were found to be highly similar between Westar and ZS11 **(Fig. 3A)**, both of which accumulated similar Na^+^ in the whole plants **(Fig. 3B)**. Further, we divided the whole plants into shoots and roots to determine whether the differences exist in the organ Na^+^ concentrations. The result showed that the root Na^+^ concentration in Westar was obviously lower than that in ZS11 **(Fig. 3C)**, which was consistent with the better root growth performance of Westar; however, the shoot Na^+^ concentration in Westar was 15.19% higher than in ZS11 **(Fig. 3C)**. Based on the shoot and root Na^+^ content **(Fig. 3D)**, we found that a higher proportion of Na^+^ retention in the roots of ZS11 than in the roots of Westar **(Fig. 3E)**. Therefore, the root-to-shoot Na^+^ translocation could not be responsible for the differential salt stress resistance between Westar and ZS11. In terms of the organ K^+^ concentrations, the shoot K^+^ concentration in Westar was obviously lower than that in ZS11 **(Fig. 3F)**; however, a reverse trend was seen in the roots, *i.e.*, the root K^+^ concentration was significantly higher in Westar than that of ZS11 **(Fig. 3F)**. It is the cytosolic Na^+^/K^+^ ratio that determines the competence of cells against salt stress **(Amin et al., 2020)**. The trends of Na^+^/K^+^ ratios in both shoots and roots were highly similar to those of Na^+^ concentrations **(Fig. 3G)**. Namely, Westar had a higher Na^+^/K^+^ ratio in the shoots and a lower Na^+^/K^+^ ratio in the roots relative to ZS11 **(Fig. 3G)**. In addition, the organ concentrations of Ca, Mg, Fe, Cu, Mn, and Zn were also evaluated to identify their roles in differential salinity resistance between Westar and ZS11. The concentrations of only root Mg^2+^and Zn^2+^ were different between the two genotypes, while the concentration profiles of the other cations were highly similar in the shoots or roots **(Supplementary Fig. S1)**. Further, we tested the concentrations of Na^+^ **(Fig. 3H)** and K^+^ **(Fig. 3I)**, and the Na^+^/K^+^ ratios **(Fig. 3J)** in the different shoot organs, including stems and different positions of leaves. Only these three parameters of the cotyledons showed significantly different levels between Westar and ZS11, while they were highly similar in the other leaves between the rapeseed genotypes **(Fig. 3H-I)**.

**Fig. 3.**
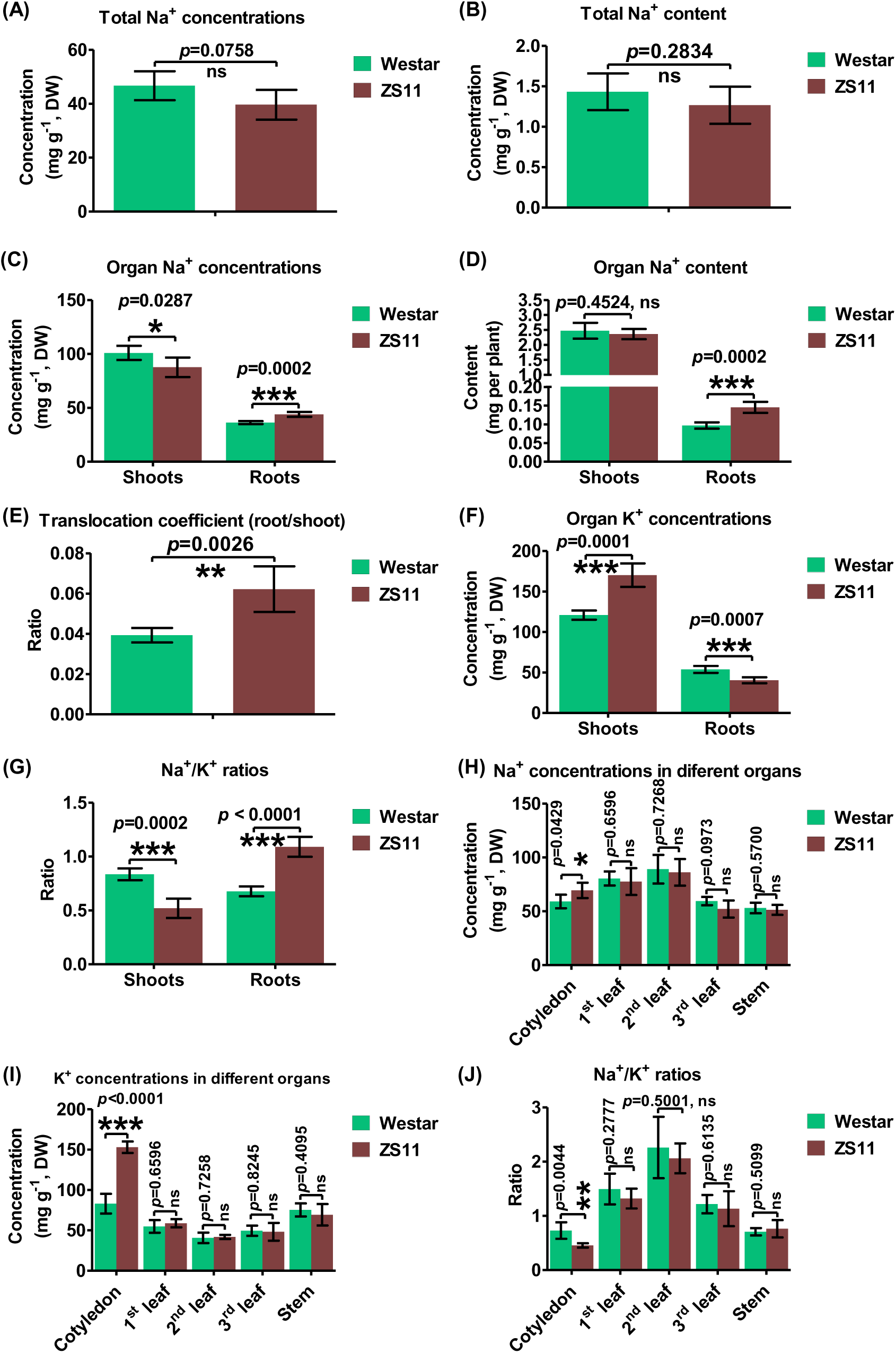
Sodium (Na) and potassium (K) ion profiling of the salt-resistant genotype Westar and the salt-sensitive genotype ZS11 under salt stress condition. **(A)** Total Na^+^ concentrations; **(B)** total Na^+^ content; **(C)** shoot and root Na^+^ concentrations; **(D)** shoot and root Na^+^ content; **(E)** Na^+^ translocation coefficient (Na^+^ content[root]/ Na^+^ content[shoot]); **(H)** total K^+^ concentrations; **(F)** shoot and root K^+^ concentrations; **(K)** shoot and root K^+^ content; **(G)** Na^+^/K^+^ ratios in the shoots and roots; **(H-J)** Na^+^ concentrations **(H)**, K^+^ concentrations **(I)**, and Na^+^/K^+^ ratios **(J)** in different shoot organs. Uniform rapeseed plants after seed germination were grown hydroponically under NaCl-free conditions for 10 d, and then transferred to 200 mM NaCl for 5 d. Ctrl, control. Data are presented as the mean (n=5) ± s.d. Significant differences (*, *p*<0.05; **, *p*<0.01; ***, *p*<0.001) were determined by unpaired two-tailed Student’s *t*-tests between two groups using the SPSS 17.0 toolkit. ns, not significant.

The ionomic profiling showed that Westar had lower Na^+^ concentrations in the roots **(Fig. 3C)**, which explained the reason for the better root growth performance of Westar than that of ZS11 under salt stress **(Fig. 2)**. The coefficient of root-to-shoot Na^+^ translocation in Westar was significantly lower than that in ZS11 **(Fig. 3E)**, and it indicated that Westar had a lower Na^+^ retention in the roots and a higher Na^+^ retention in the shoots. To further identify whether vacuolar Na^+^ compartmentation was implicated in the salt stress resistance in the roots, TEM coupled with X-ray energy spectrum analysis was used to determine the root flux subcellular redistribution and of Na^+^ under salt stress **(Fig. 4A-D)**. The result showed that the relative Na^+^ abundances did not present obvious differences in neither the meristematic zones nor the elongation zones between the roots of Westar and ZS11 **(Fig. 4D)**. Then, we used the NMT to test the differences in the root Na^+^ extrusion, and it remained to be highly similar between Westar and ZS11 **(Fig. 4E-G)**, the root K^+^ influx rate of which was significantly lower than that of Westar **(Fig. 4E, H, I)**.

**Fig. 4.**
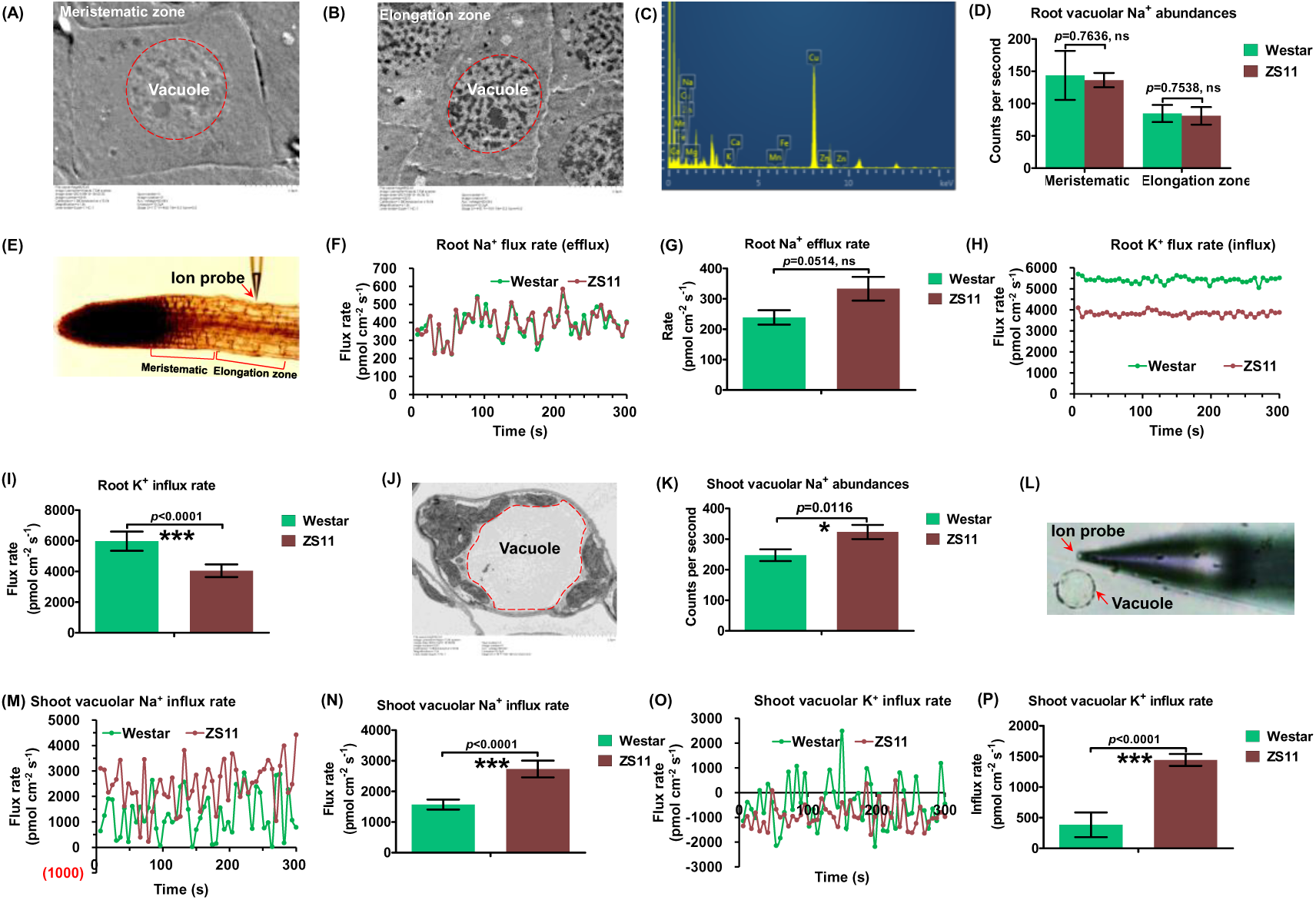
Non-invasive micro-test technology and transmission electron microscopy coupled with X-ray analysis for the determination of root flux and subcellular redistribution of Na^+^/K^+^ under salt stress. **(A-B)** TEM- assisted ultrastructure identification of the root cells in the meristematic **(A)** and elongation **(B)** zones of rapeseed plants under salt stress. **(C)** X ray spectrum for the identification of ion abundances. **(D)** X ray spectrum- assisted analysis of relative Na^+^ abundances in the root cells in the meristematic and elongation zones of Westar and ZS11 under salt stress. (E) Representative measurement graph of Na^+^/K^+^ flux in the cells in the meristematic and elongation zones of rapeseed roots under salt stress. **(F-I)** Flux kinetics **(F, H)** and net flux rate **(G, I)** of Na^+^ **(F, G)** and K^+^ **(H, I)** in the roots of Westar and ZS11 under salt stress. **(J)** TEM-assisted ultrastructure identification of the shoot cells of rapeseed plants under salt stress. **(K)** X ray spectrum-assisted analysis of relative Na^+^ abundances in the shoot vacuoles of Westar and ZS11 under salt stress. **(L)** Representative measurement graph of Na^+^/K^+^ flux in the shoot vacuoles of rapeseed plants under salt stress. **(M-P)** Flux kinetics **(M, O)** and net flux rate **(N, P)** of Na^+^ **(M, N)** and K^+^ **(O, P)** in the shoot vacuoles of Westar and ZS11 under salt stress. Uniform rapeseed plants after seed germination were grown hydroponically under NaCl-free conditions for 10 d, and then transferred to 200 mM NaCl for 5 d. Ctrl, control. Data are presented as the mean (n=5) ± s.d. Significant differences (*, *p*<0.05; **, *p*<0.01; ***, *p*<0.001) were determined by unpaired two-tailed Student’s *t*-tests between two groups using the SPSS 17.0 toolkit. ns, not significant.

The result aforementioned revealed that Westar had a higher shoot Na^+^ concentration **(Fig. 3C)**, while it was more resistant to salt stress compared with ZS11 **(Fig. 1)**. It indicated that a more complex mechanism might be implicated in the regulation of differential salt stress resistance in the shoots than in the roots between Westar and ZS11. Vacuolar Na^+^ compartmentation confers resistance against salt stress in crop species (İ**brahimova et al., 2021)**. First, we investigated the relative Na^+^ abundances in the shoot vacuoles of Westar and ZS11 through TEM coupled with X-ray energy spectrum analysis **(Fig. 4J)**; the result showed that ZS11 had a significantly higher vacuolar Na^+^ abundance in the shoots than Westar **(Fig. 4K)**. Then, the analysis of NMT-assisted ion kinetics showed that ZS11 had higher influx rates of both Na^+^ and K^+^ than Westar **(Fig. 4L-P)**. Taken together, it could be concluded that shoot vacuolar Na^+^ sequestration was not involved in the differential salt stress resistance between Westar and ZS11.

Previous studies showed that ZS11 is more resistant against boron deficiency and cadmium toxicity than Westar; moreover, the differential salt resistance between the two rapeseed genotypes is closely correlated with their cell wall characteristics **(Hua et al., 2016; Zhang et al., 2019)**. Currently, more and more studies revealed that cell wall plays a pivotal role in the resistance of plants against salt stress **(Dabravolski and Isayenkov, 2023)**. Therefore, it promoted us to investigate the differences in the cell wall Na^+^ retention between Westar and ZS11. Under salt stress, the chloroplasts became more severely deformed in ZS11 than in Westar **(Fig. 5A, B)**. Salt stress may cause cell wall softening, notably through its effect on pectins **(Feng et al., 2018)**. Cell ultrastructure analysis showed that the cell wall of ZS11 was much thicker than that of Westar under salt stress **(Fig. 5C, D)**. Further, we investigated the Na^+^ concentrations of different cell wall components in the shoots of Westar and ZS11, and the Na^+^ concentration in the cell wall of Westar was almost two folds of that in the cell wall of ZS11 **(Fig. 5E)**. The concentrations of Ca, Mg, Mn, and Cu in the cell walls of ZS11 were significantly higher than those of Westar, while the concentrations of Fe and Zn did not present obvious difference between the rapeseed genotypes **(Supplementary Fig. S2)**. The pectin Na^+^ concentration of Westar was up to 8 folds that of ZS11 **(Fig. 5F)**, although the Na^+^ concentrations were also significantly different between Westar and ZS11 in the cell walls minus pectins **(Fig. 5G)**. Based on the Na^+^ profiling in the cell wall components, we found that the concentration ratio of Na^+^ /Na^+^ in Westar was significantly higher than that in ZS11 **(Fig. 5H)**, which indicated that the pectin of Westar might have a stronger ability of Na^+^ retention in the cell wall than that of ZS11. Both the K^+^ concentrations **(Fig. 5I-K)** and the Na^+^/K^+^ ratios **(Fig. 5L)** in total cell walls, pectins, and the other cell wall components in Westar were significantly higher than those in ZS11 **(Fig. 5I-L)**. Comparative analysis of cell wall components showed that the abundances of cellulose **(Fig. 5M)**, lignin **(Fig. 5N)**, and pectins (including covalently/ionically binding pectin) **(Fig. 5O, P)** did not show obvious differences between Westar and ZS11 **(Fig. 5M-P)**. Therefore, we further tested the modification status of cell walls and pectins, and found that both the absorbance of cell wall and the relative abundance of carboxyl groups in Westar were markedly higher than those of ZS11 **(Fig. 5Q, R)**. Methylation degree analysis showed that the pectins of Westar had a higher methylation degree than that of ZS11 **(Fig. 5S)**, which was then confirmed to have a lower PME activity compared with Westar **(Fig. 5T)**. Taken together, pectin methylation-mediated cell wall Na^+^ retention might play a very important role in the regulation of differential salt stress resistance between Westar and ZS11.

**Fig. 5.**
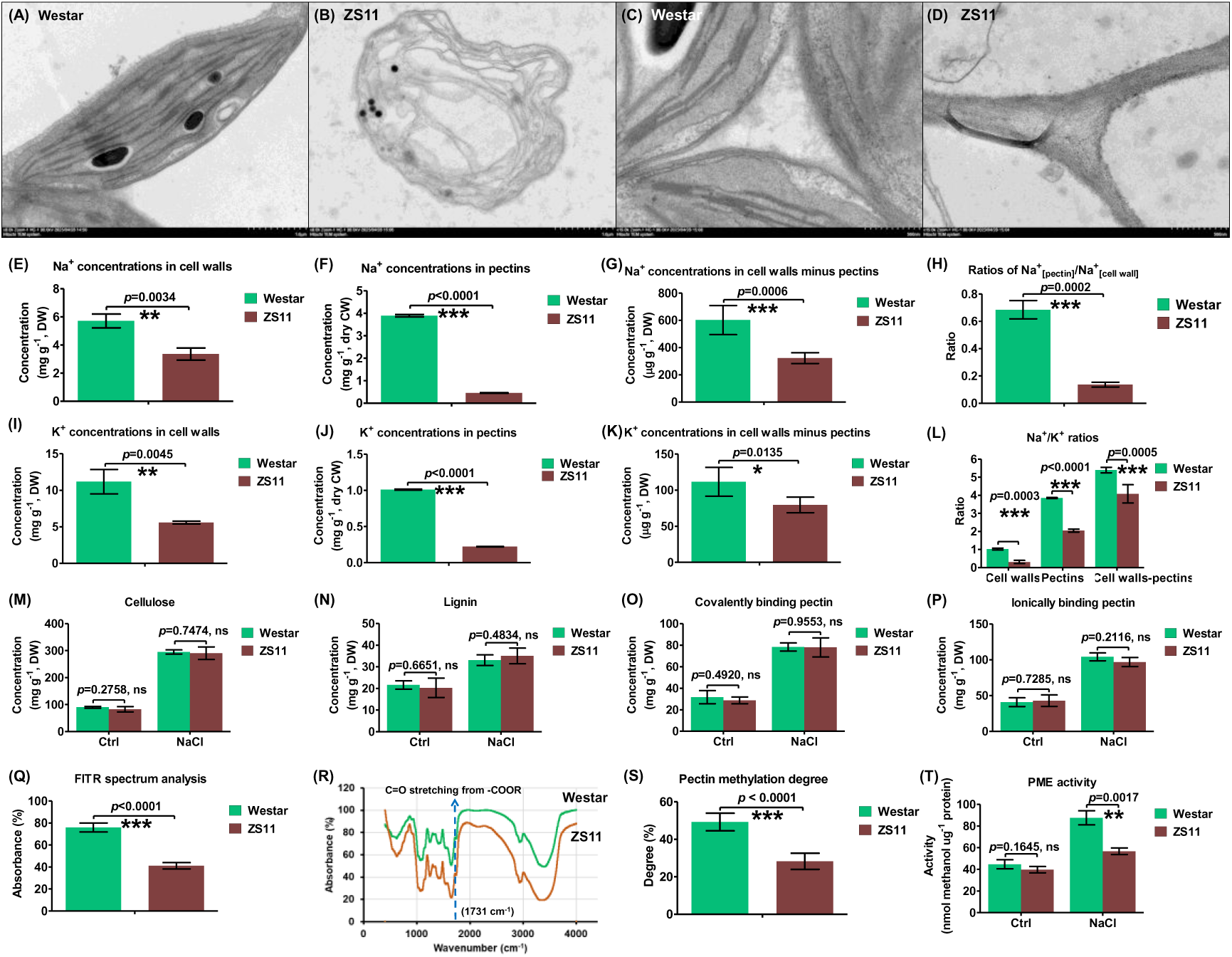
Comparative analysis of components and Na^+^/K^+^ profiling in the cell walls of the salt-resistant genotype Westar and the salt-sensitive genotype ZS11 under control and salt stress conditions. **(A-D)** Chloroplast **(A, B)** and cell wall **(C, D)** ultrastructure of Westar **(A, C)** and ZS11 **(B, D)** under salt stress. Scale bar = 1 μm. **(E-K)** Concentrations of Na^+^ and K^+^ in the cell walls **(E, I)**, pectins **(F, J)**, and cell walls minus pectins **(G, K)** in the shoots of Westar and ZS11 under salt stress. **(H)** Ratios of pectin Na^+^ concentrations/cell wall Na^+^ concentrations. **(L)** Na^+^/K^+^ ratios in the cell wall, pectins, and cell walls minus pectins in the shoots of Westar and ZS11 under salt stress. **(M-P)** Concentrations of cellulose **(M)**, lignin **(N)**, covalently binding pectin **(O)**, and ionically binding pectin **(P)** in the shoots of Westar and ZS11 under control and salt stress. **(Q)** Relative abundances of carboxyl groups (COO^−^) in cell walls. **(R)** Fourier transform infrared spectral analysis of the cell walls at the wavenumber ranging from 4000 to 400 cm^-1^. **(S)** Methylation degree of the pectins isolated from shoot cell walls. **(T)** Activity of pectin methylesterase (PME) in the shoots of Westar and ZS11 under control and salt stress. Uniform rapeseed plants after seed germination were grown hydroponically under NaCl-free conditions for 10 d, and then transferred to 200 mM NaCl for 5 d until sampling. Ctrl, control. Data are presented as the mean (n=5) ± s.d. Significant differences (*, *p*<0.05; **, *p*<0.01; ***, *p*<0.001) were determined by unpaired two-tailed Student’s *t*-tests between two groups using the SPSS 17.0 toolkit. ns, not significant.

### Identification and characterization of genome-wide DNA variations between Westar and ZS11

To identify the genome-wide DNA variations between the salt-resistant genotype Westar and the salt-sensitive genotype ZS11, we used their public genome sequences to compare their genomic polymorphisms. According to the current genome assembly, the genome sizes of Westar and ZS11 were up to 1,007 and 1,008 Mb, being annotated with 97,514 and 100,919 genes, respectively **(Song et al., 2020)**. In general, the genome of Westar had a good collinearity with that of ZS11 (**Fig. 6A, Supplementary Fig. S3**); as shown in **Fig. 6B**, thousands of genomic variants, including 2,429,398 SNPs, 772,724 InDels, 36,912 SVs, and 103,457 CNVs, between Westar and ZS11, were non-evenly distributed across the 19 chromosomes (A1-A10, C1-C9) of *B. napus* **(Supplementary Figs. S4, 5)**.

**Fig. 6.**
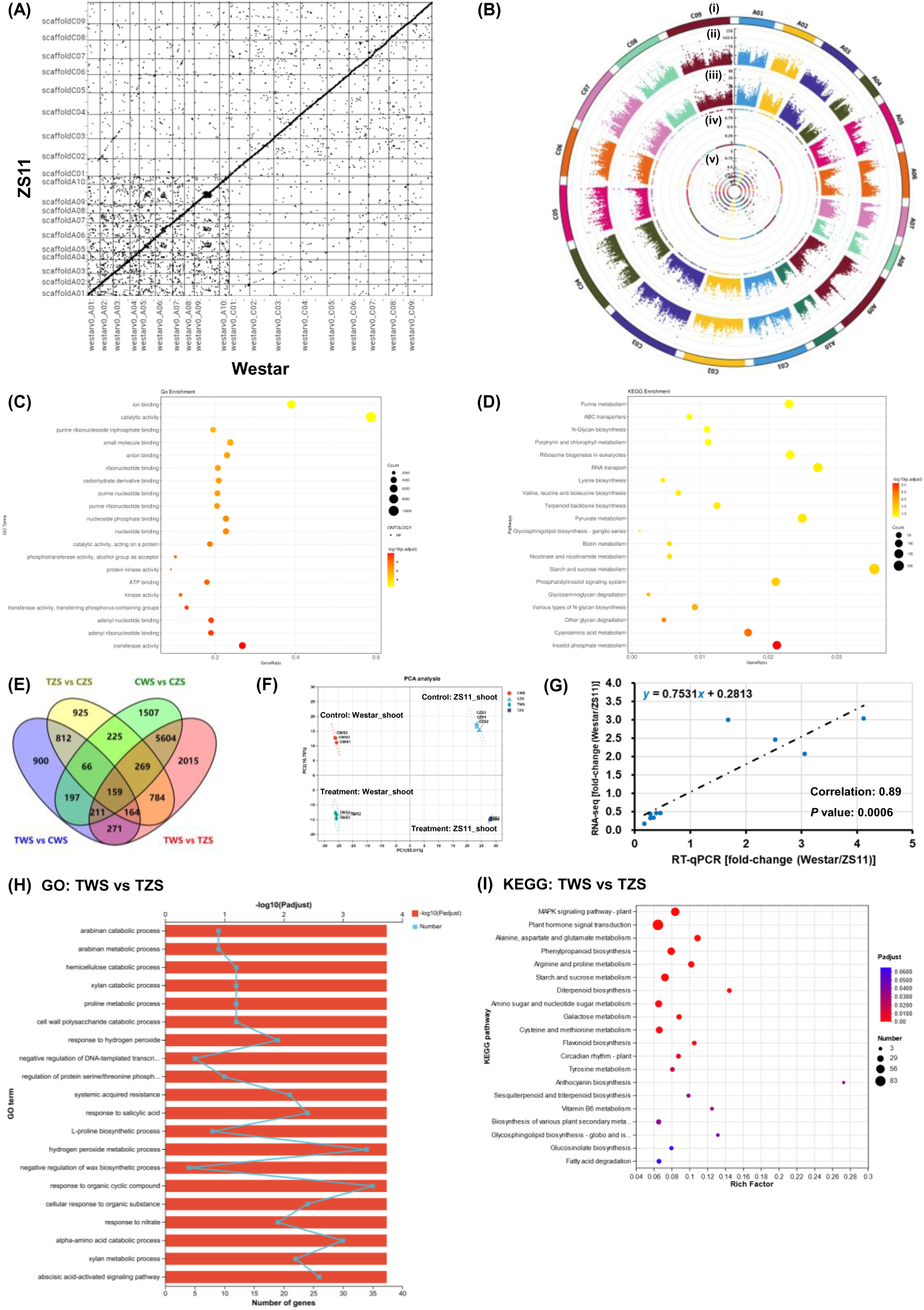
Overview of the genomic DNA polymorphisms and genome-wide differentially expressed genes between the salt-resistant genotype Westar and the salt-sensitive genotype ZS11 under control and salt stress conditions. **(A)** Comparative analysis of genome synteny between Westar and ZS11. **(B)** Overview of the genome-wide genetic variants between Westar and ZS11, which are delineated by the Circos program. In the Circos figure, the variants are as follows outside-to-inside: chromosomes (i), single nucleotide polymorphisms (SNPs) (ii), insertions/deletions (InDels) (iii), copy number variation (CNV, iv), structure variations (SVs, v). **(C, D)** Enrichment analysis of gene ontology/GO **(C)** and Kyoto Encyclopedia of Genes and Genomes/KEGG **(D)** terms for the genes detected with DNA polymorphisms. **(E)** Venn diagrams for the genome-wide differential expressed genes (DEGs) in the shoots (S) between Westar (W) and ZS11 (Z) under control and salt stress conditions. C, control (NaCl-free); T, treatment (200 mM NaCl). **(F)** Principal component analysis (PCA) of the genome-wide DEGs. **(G)** Consistency analysis between the RT-qPCR assays and RNA-seq results. **(H- I)** GO **(H)** and KEGG **(I)** enrichment analysis of the DEGs between Westar and ZS11 under salt stress. For the transcriptome sequencing, uniform rapeseed plants after seed germination were grown hydroponically under NaCl-free conditions for 10 d, and then transferred to 200 mM NaCl for 3 d until sampling.

In total, the genome-wide SNPs ranged from 157,429 (Chr. A3) to 39,822 (Chr. A8) (**Supplementary Fig. S5A)**, and the nucleotide diversity π (average SNP number per nucleotide) ranged from 1.18 × 10^−3^ (Chr. C1) to 4.53 × 10^−3^ (Chr. A4), with an average value of 3.21 × 10 ^− 3^ and 2.16 × 10 ^− 3^ on the A and C subgenomes, respectively (**Supplementary Fig. S5B)**. The annotations of reference (ZS11) genome were used to examine the distribution of SNPs within various genomic features. The detected SNPs were categorized into two groups: transitions (A/G and C/T; Ts) and trans-versions (A/C, A/T, C/G and G/T; Tv) based on the nucleotide variations between Westar and ZS11. Among the genome-wide SNPs, 57.20% belonged to the transition type, which was more than that of the transversions **(Supplementary Fig. S6A)**. The ratio of Ts to Tv was approximately 1.34, which was larger than the expected value (0.5). The intergenic SNPs accounted for the largest proportion (46.80%) of the genome-wide SNPs **(Supplementary Fig. S6B)**; among the SNPs located in the genic areas, 45.98% of the variations occurred in the coding sequences **(Supplementary Fig. S6B)**. Only a minor fraction (0.56%) of the SNPs located in the exonic regions were mapped onto the stop codons **(Supplementary Fig. S6C)**. In general, the ratio of synonymous SNPs (59.57%) to non-synonymous SNPs (39.88%) was about 1.49 **(Supplementary Fig. S6C)**. Also, the genome-wide 772,724 InDels ranged from 17,108 (Chr. C6) to 59,076 (Chr. C3) with an average of 18,250 InDels on each chromosome **(Supplementary Fig. S5C)**. The majority of (45.71%) total InDel variants occurred in the intergenic regions, whereas 15.94% of InDels were identified in the genic sequences **(Supplementary Fig. S6E)**. Among the InDels occurring in the exons, only ∼7.43% were identified in the stop codons **(Supplementary Fig. S6F)**. For the other InDels, a total of 43.23% caused frame-shift deletion or insertion **(Supplementary Fig. S6F)**. In total, the genome-wide SVs ranged from 2,968 (Chr. A3) to 795 (Chr. A8) (**Supplementary Fig. S5D).** The types of SVs between Westar and ZS11 were divided into five terms: genomic fragment insertion, deletion, intrachromosomal translocation, and inter-chromosomal translocation. As expected, most SVs led to inter-chromosomal translocation, deletion, followed by genomic fragment deletion, whereas genomic insertion was the least **(Supplementary Fig. S7A-D)**. In total, the genome-wide CNVs ranged from 71 (Chr. C3) to 7 (Chr. A8) (**Supplementary Fig. S5E)**, and the CNVs contain two types: duplication and deletion, and copy number deletions (1,677) of genes were similar to gene duplications (1,699) between Westar and ZS11 **(Supplementary Fig. S8)**.

GO enrichment analysis of the genome-wide DNA variants between Westar and ZS11 were highly enriched in the categories of ion binding, catalytic activity, and transferase activity **(Fig. 6C)**, while the KEGG analysis revealed that the pathways involving carbohydrate (starch and sucrose) metabolism were over-accumulated **(Fig. 6D)**.

### Genome-wide differential expression profiling between Westar and ZS11 under salt stress

To identify the core DEGs regulating differential salt stress resistance between Westar and ZS11, a high-throughput comparative genome-wide transcriptome sequencing was performed for the two rapeseed genotypes. After removing adapter sequences and low-quality reads, approximately 5.8 × 10^7^ clean reads were obtained from each sample. The total length of clean reads from the 24 samples reached about 1.01 × e^11^ nt with Q_20_ > 98% and Q_30_ > 94% **(Supplementary Table S2)**. Most of *Pearson* correlation coefficients between each pair of biological replicates of each rapeseed genotype under the same treatment were > than 0.90 **(Supplementary Fig. S9)**. Clustering trees, presenting the distances among biological replicates, showed comparable heights among the three biological replicates of each sample. Moreover, the hierarchical clustering of genome-wide gene expression revealed that expression patterns were similar among the three biological replicates of each sample **(Supplementary Fig. S10)**. Taken together, these parameters indicated that the transcriptome sequencing data were of high quality among different biological replicates.

In the shoots, a total of 2,780 and 3,404 salt-responsive genes were identified in Westar and ZS11, respectively **(Fig. 6E, Supplementary Table S3)** including 164 salt-responsive DEGs between them **(Fig. 6E)**. Principal component analysis showed that different rapeseed genotypes or treatments exhibited significantly different transcriptomic features in terms of the control or salt stress conditions **(Fig. 6F)**. Ten DEGs **(Supplementary Table S4)** were randomly selected to compare their expression correlations between the RT-qPCR assays and transcriptome sequencing. The results showed that the gene expression patterns using the two assay methods highly correlated (correlation coefficient, 0.89; *p* value, 0.0006) **(Fig. 6G)**.

GO enrichment analysis of the genome-wide DEGs between Westar and ZS11 revealed that under control condition, the terms of organic substance biosynthesis and cellular biosynthesis were highly accumulated **(Supplementary Fig. S11A)**, while the categories of systematic acquired resistance, response to salicylic acid, hydrogen peroxide metabolic process, xylan metabolic process, and abscisic acid-activated signaling pathway were over-presented under salt stress **(Fig. 6H)**. KEGG pathway analysis showed that under control condition, the metabolism of phosphonate/ phosphinate, sphingolipid/glycosphingolipid, and glycosaminoglycan was highly enriched **(Supplementary Fig. S11B)**, while the terms of plant hormone signal transduction, proline metabolism, and anthocyanin biosynthesis were over-accumulated under salt stress **(Fig. 6H)**.

### Genome-wide identification and multi-omics assisted molecular characterization of the core *PME* member(s) regulating differential salt stress resistance between Westar and ZS11

First, we employed the transcriptomic data to investigate the transcriptional profiling of some potential Na^+^ transporters, which are responsible for cellular Na^+^ influx, compartmentation, and extrusion **(Fig. 7A)**. As shown in **Fig. 4**, we found that the salt-sensitive genotype ZS11 had stronger vacuolar Na^+^ sequestration than the salt-resistant genotype Westar; consistently, we identified only the expression of *BnaC5.NHX3* and *BnaA10.NHX4* was significantly different between Westar and ZS11, which had much higher expression levels of these two genes (Fig. 7B).

**Fig. 7.**
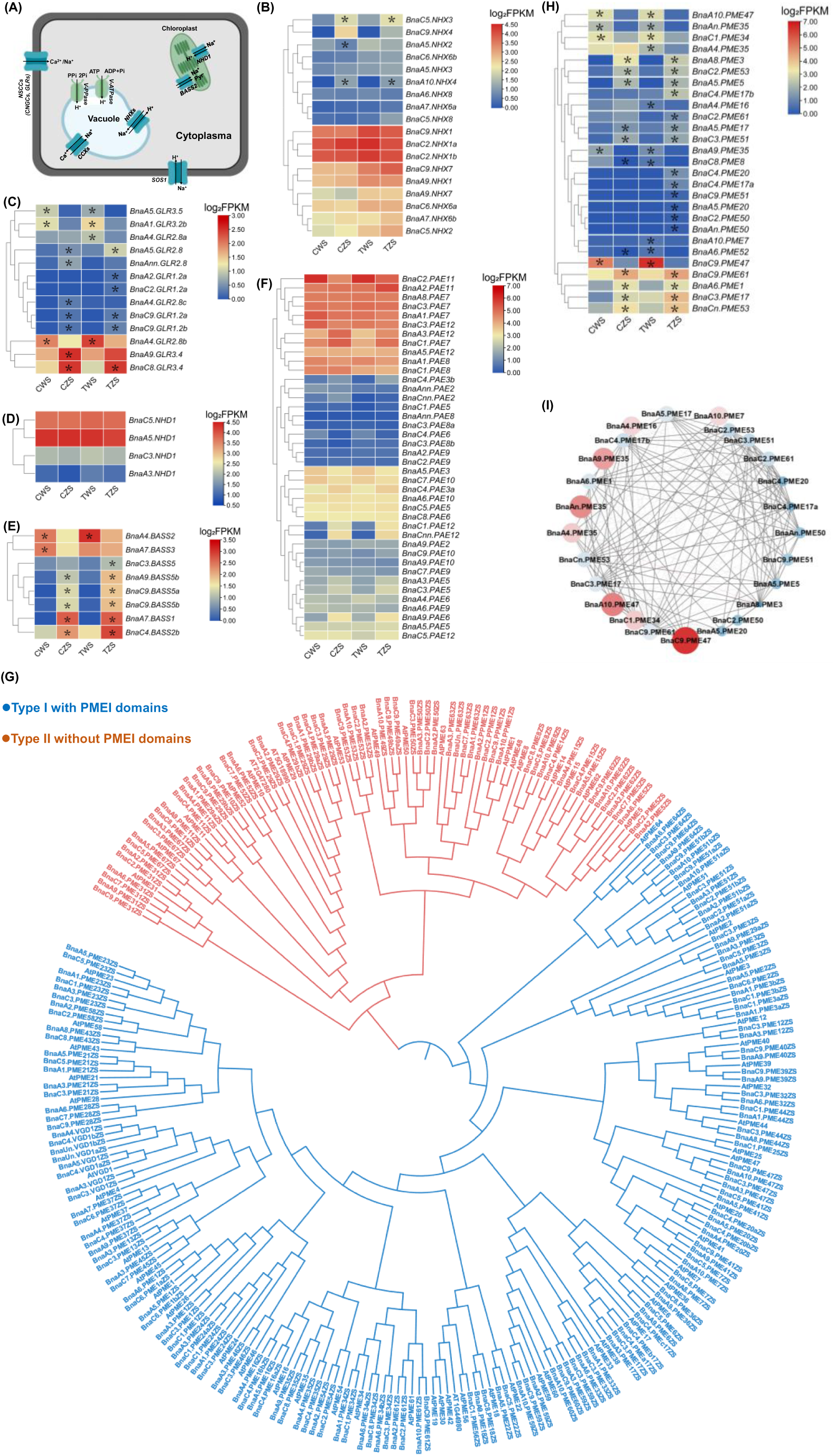
Transcriptional profiling of the cellular Na^+^ transporter genes and genome-wide identification and transcriptional characterization of *pectin methylesterase* (*PME*) family genes. **(A)** A molecular model showing the genes involved in the transport of Na^+^. BASS, bile acid sodium symporter; CNGC, cyclic nucleotide-gated channel; GLR, glutamate-like receptor; NHX/NHD, Na^+^/H^+^ antiporter; SOS, salt overly sensitive. **(B-F)** Transcriptional profiling of *NHXs* **(B)**, *GLRs* **(C)**, *NHDs* **(D)**, and BASSs **(E)**, and *PAEs* **(F)**. PAE, pectin acetylesterase. **(G)** Phylogeny analysis of the *PMEs* in Arabidopsis (*Arabidopsis thaliana* L.) and rapeseed (*Brassica napus* L., cv. ZS11). The *PMEs* indicated by blue and red color belong to the type I and II members, respectively. **(H)** Differential expression profiling of the *PMEs* in the shoots (S) between Westar (W) and ZS11 (Z) under control and salt stress conditions. **(I)** Co-expression analysis of the differentially expressed PMEs between Westar and ZS11. C, control (NaCl-free); T, treatment (200 mM NaCl). The genes with an absolute log2[fold change(Westar/ZS11)] > 1.0 and an adjusted *p* value < 0.05 were considered to be differentially expressed. The DEGs with higher expression between the salt-resistant genotype Westar and the salt-sensitive genotype ZS11 are denoted by asterisks. The heatmaps show gene expression levels as indicated by log2FPKM values (fragments per kilobase of exon model per million mapped reads). The gene co-expression networks were constructed using Cytoscape (http://www.cytoscape.org/). Cycle nodes represent genes, and the size of the nodes represents the power of the interrelation among the nodes by degree value. Edges between two nodes represent *PME* interactions.

PMEs are classified based on the presence or absence of the PMEI domain. The type I of PMEs has 1-3 PMEI domains, while type II has no PMEI domain **(Kanneganti and Gupta, 2009)**. There are 66 *PMEs* annotated in the Arabidopsis genome **(Louvet et al., 2006)**; in this study, through genome-wide retrieval of the rapeseed *PME* family genes, a total of 209 (including 141 type I and 68 type II) and 180 (including 129 type I and 51 type II) members, unevenly distributed across the rapeseed chromosomes **(Supplementary Figs. S12, 13)**, were characterized in the Westar and ZS11 genome, respectively **(Supplementary Table S5)**. The Ka/Ks values of all *BnaPMEs* were obviously smaller than 1.0 in both Westar and ZS11 **(Supplementary Table S5)**, which indicated that *BnaPMEs* underwent strong purifying/negative selection. Phylogenetic relationship analysis showed that the type I and type II of genome-wide *BnaPMEs* were obviously clustered into two clades **(Fig. 7G; Supplementary Fig. S14A)**. Further synteny analysis showed that a large number of inversion of *BnaPMEs* occurred between Westar and ZS11 **(Supplementary Fig. S14B)**.

Among the genome-wide *PMEs*, there were 27 members differentially expressed in the shoots between Westar and ZS11 **(Fig. 7H)**. Genome-wide expression profiling of *BnaPMEs* and gene co-expression network analysis showed that *BnaC9.PME47*, almost with the highest expression abundances **(Supplementary Fig. S15)**, might play a core role in the *PME* family **(Fig. 7I)**. The above phylogeny analysis showed that BnaC9.PME47, similar to AtPME47, belonged to the type I without PMEI domains **(Fig. 7G)**. In turn, the RNA-seq and RT-qPCR results showed that the *BnaC9.PME47* expression was significantly induced by salt stress **(Figs. 6G, 7H)**.

Through a genome-wide association study, a QTL for salt stress resistance at the end (about 62.56 Mb) of the chromosome C9 was detected in rapeseed **(Yong et al., 2015; Fig. 8A)**, which was also identified in another previous study **(Lang et al., 2017)**. *BnaC9.PME47* was consistently located in the two reported QTL regions for salt stress resistance in rapeseed **(Fig. 8C)**, which confirmed the pivotal role of *BnaC9.PME47* in conferring rapeseed salt stress resistance. All of *AtPME47*, *BnaC9.PME47*^Westar^, and *BnaC9.PME47*^ZS11^ had similar genic organization, including four exons and three introns, although they presented different genomic lengths **(Fig. 8D)**. Higher plant PMEs are often pre-pro-proteins, in which the mature, active part of the protein (PME domain, IPR000070) is preceded by an N-terminal extension (PRO region) that varies in length and amino acid identity between isoforms. The PRO region is similar to the PME inhibitor domain (PMEI domain, IPR006501). *BnaC9.PME47*^Westar^ and *BnaC9.PME47*^ZS11^ had two amino acid variations at the 96^th^ and 234^th^ (located in the PMEI domain) residue position **(Fig. 8E)**. Phylogeny analysis showed that the rapeseed *BnaC9.PME47* was derived from its diploid ancestor *BolC9.PME47* **(Fig. 8F)**. Molecular characterization of tissue-specific expression patterns revealed that *BnaC9.PME47* was preferentially expressed in the shoots, including leaves **(Fig. 8G)**. In terms of the differential expression of *BnaC9.PME47* between Westar and ZS11, we investigated the differences in the promoter region sequences, and found 18 SNPs and seven InDels between the rapeseed genotypes **(Supplementary Fig. S16)**. Analysis of *cis*-acting regulatory elements showed that there were four annotated elements around these genomic polymorphic sites, among which the ABA responsive element was most noteworthy **(Supplementary Table S6)**. Further, natural variation of the promoter region sequences of *BnaC9.PME47* in eight rapeseed genotypes, whose genome information has been publicly released in the BnIR database, showed that they were clustered into two haplotypes: the type I only contained Westar, while the other seven rapeseed genotypes belonging to the type II had same sequences as ZS11 **(Fig. 8H)**. Moreover, Westar outperformed than the other seven rapeseed genotypes under salt stress **(Fig. 8I)**, and the expression of *BnaC9.PME47* in Westar was significantly higher than that in the other rapeseed genotypes **(Fig. 8J)**.

**Fig. 8.**
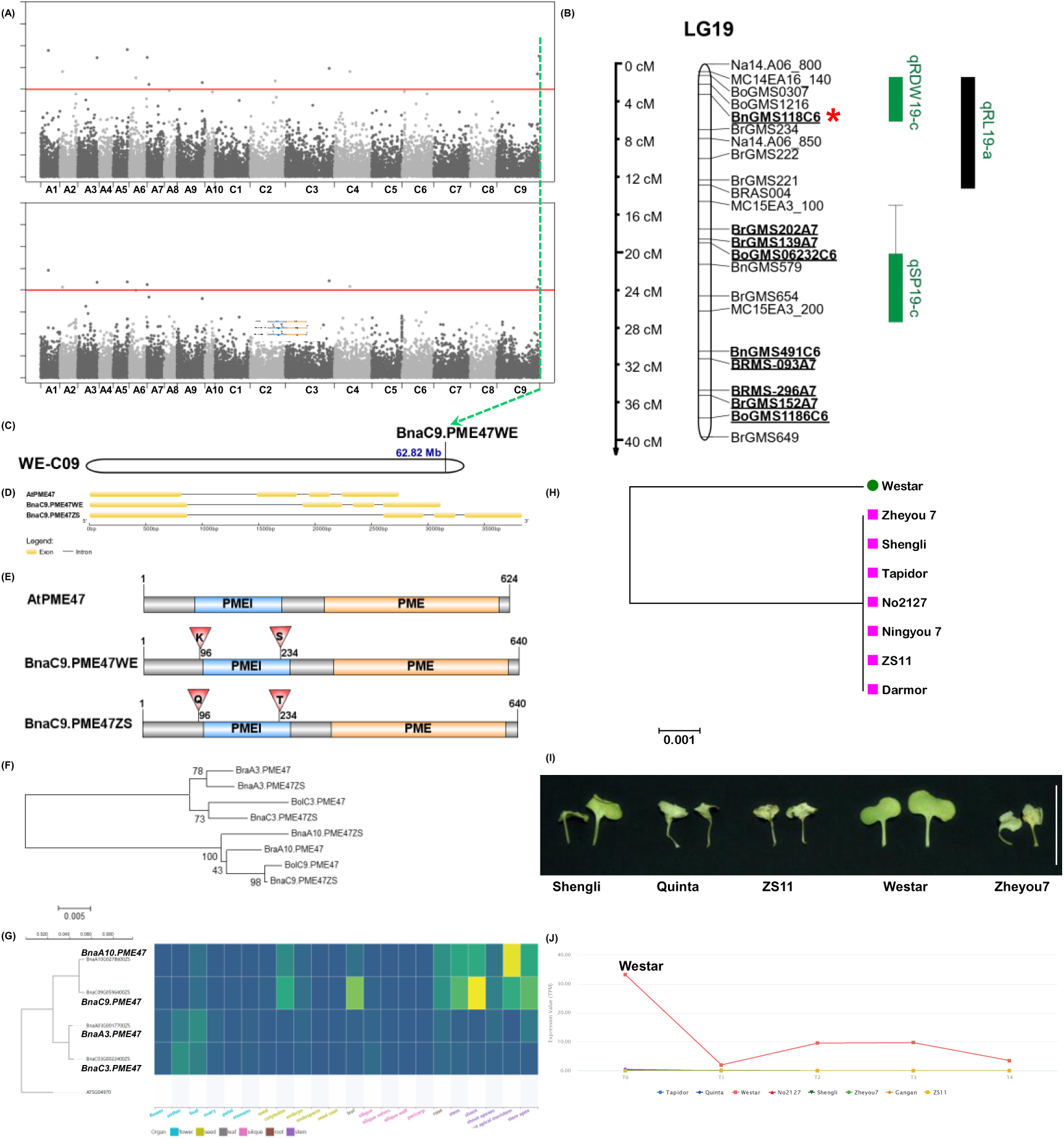
Natural variation of *BnaC9.PME47* in rapeseed genotypes. **(A-B)** Two mapped genetic loci controlling rapeseed salt stress resistance previously reported by genome-wide association analysis **(A, Yong et al., 2015)** and quantitative trait locus (QTL) mapping **(B, Lang et al., 2017)**. **(C-G)** Physical position **(C)**, gene structure **(D)**, amino acid sequence variation **(E)** of *BnaC9.PME47* in *B. napus*. The position of *BnaC9.PME47* in the mapped genomic regions is labelled by crashed lines in **Fig. 7A** and by an asterisk in **Fig. 7B**. **(F)** Phylogenetic analysis of the PME47 proteins in *Brassica rapa*, *Brassica oleracea*, and *Brassica napus* (cv. ZS11). (G) tissue-specific expression characteristics in various organs of rapeseed plants. **(H)** Phylogenetic analysis of the promoter sequences of *BnaC9.PME47* in rapeseed genotypes. The evolutionary history was inferred using the neighbor-joining method. The scale bar corresponds to a distance of 0.01 substitutions per site. Bootstrap values are indicated adjacent to the corresponding node. **(I)** Growth performance of different rapeseed genotypes under salt stress. Scale bar: 5 cm. Uniform rapeseed plants after seed germination were grown hydroponically under NaCl-free conditions for 10 d, and then transferred to 200 mM NaCl for 5 d until sampling. **(J)** Comparative analysis of the *BnaC9.PME47* expression, downloaded from the BnIR database (https://yanglab.hzau.edu.cn/BnIR), in different rapeseed genotypes. T0: 24 days after sowing; T1: 54 days after sowing; T2: 82 days after sowing; T3: 115 days after sowing; T4: 147 days after sowing.

To further identify the function of *BnaC9.PME47* in regulating salt stress resistance, we tested its subcellular localization in the epidermal cells of tobacco leaves. Before plasmolysis, we could not identify BnaC9.PME47 was localized on the plasma membrane or the cell wall **(Fig. 9A)**; however, after plasmolysis, both BnaC9.PME47^Westar^ and BnaC9.PME47^ZS11^ were obviously localized on the cell wall, dissociated from the plasma membrane **(Fig. 9A, B)**. Considering that the function of *PME47* has been not identified and characterized in the model Arabidopsis plants, we used two homozygous *AtPME47* T-DNA insertion mutants (*atpme47-1* and *atpme47-2*) **(Supplementary Fig. S17)** to identify their growth performance compared with the wild type (Col-0) under salt stress. T-DNA was inserted into an intron and the promoter region in the *atpme47-1* and *atpme47-2* mutants, respectively **(Fig. 9C)**. Under control condition, the two T-DNA insertion mutants of *AtPME47* and the wild type grew similarly; while under salt stress, only *atpme47-2* showed obvious leaf chlorosis than the wild type **(Fig. 9D)**. Considering that the T-DNA insertion into the intron of *AtPME47* did not cause the alteration of growth performance in *atpme47-1*, we only investigated the physiological parameters of *atpme47-2* and wild type in the following experiments. Under salt stress, the root system architecture of *atpme47-2* was much weaker than that of the wild type although they had similar performance under control condition **(Fig. 9E)**. In detail, the total root length and root tip number of *atpme47-2* was significantly lower than those of the wild type under salt stress **(Fig. 9F, G)**. Further ionomic profiling showed that although the Na^+^ concentrations in the whole plants of *atpme47-2* were significantly smaller than those of the wild type, *atpme47-2* showed higher sensitivity to salt stress than the wild type **(Fig. 9H)**. In addition, the concentrations of several essential metal ions, including K, Ca, Mg, Fe, Cu, Mn, and Zn, in the whole plants of *atpme47-2* were also significantly smaller than those of the wild type **(Fig. 9H)**.

**Fig. 9.**
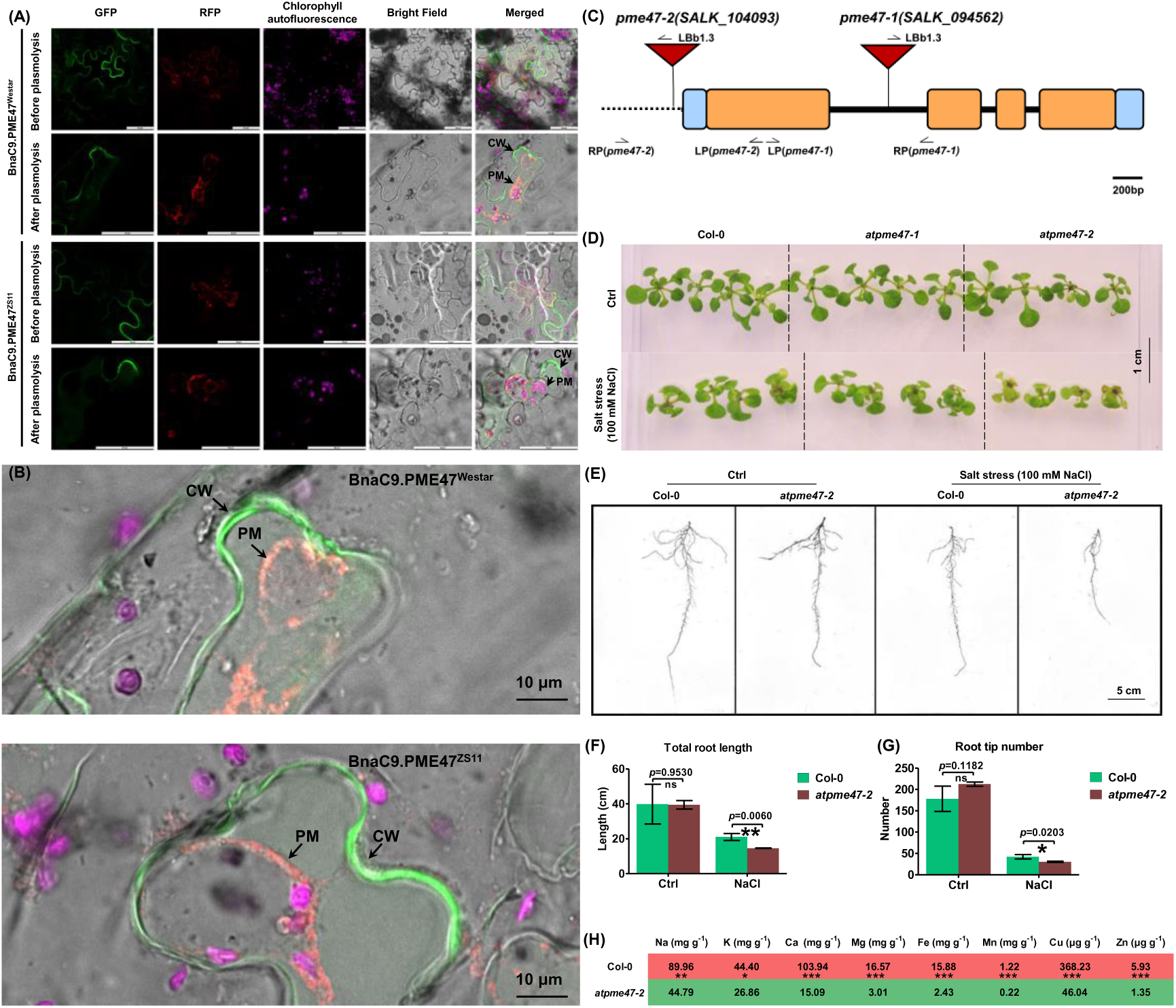
Subcellular localization of *BnaC9.PME47* and identification of phenotype performance of *AtPME47* T-DNA mutants under salt stress. (A-B) Low-magnification **(A)** and close-up **(B)** view of subcellular localization of *BnaC9.PME47*^Westar^ and *BnaC9.PME47*^ZS11^ in epidermal cells of tobacco leaves. CW, cell wall. PM, plasma membrane. In **Fig. 9A**, scale bar: 50 μm; In **Fig. 9B**, scale bar: 10 μm. **(C)** Schematic diagram of the *AtPME47* T-DNA insertion mutants, including *atpme47-1* (SALK_104093) and *atpme47-2* (SALK_094562). Untranslated regions, exons, and introns are indicated by blue boxes, orange boxes, and black lines, respectively. Primers used in the genotype analysis are indicated by arrows. **(D)** Shoot growth performance of wild type (Col-0) and the *atpme47* T-DNA insertion mutants (*atpme47-1* and *atpme47-2*) under control and salt stress conditions. Scale bar: 1 cm. **(E-H)** Root growth performance **(E)**, total root length **(F)**, root tip number **(G)**, and ion profiling **(H)** of the wild type (Col-0) and *atpme47-2* T-DNA insertion mutants under control and salt stress conditions. Scale bar: 5 cm.

### Differential phytohormone profiling between Westar and ZS11 under salt stress

The transcriptomic results showed that the GO and KEGG analysis of the differentially expressed genes between Westar and ZS11 under salt stress showed that response to salicylic acid, abscisic acid-activated signaling pathway, and plant hormone signal transduction were over-accumulated under salt stress **(Fig. 6H, I)**. It indicated that phytohormones might play key roles in the cell wall Na^+^ retention mediated differential responses between Westar and ZS11 to salt stress. The concentrations of both IAA and IBA, key types of endogenous auxin in plants, were highly similar in the shoots between Westar and ZS11, which had significantly higher IBA concentrations in the roots than Westar **(Fig. 10A, B)**. We tested six types of cytokinin, including *c*Z, *t*Z, iP, iPA, *t*ZR, and KT. The results showed that the concentrations of *c*Z, *t*Z, and iPA were significantly higher in the shoots of ZS11 than in the shoots of Westar **(Fig. 10C-H)**, which had a much higher level of *t*ZR than ZS11 in the roots **(Fig. 10E)**. In terms of GA, all the concentrations of shoot GA_7_ and root GA_3_ and GA_7_ presented higher levels in ZS11 than in Westar **(Fig. 10I-K)**. In the shoots, all the concentrations of ABA, JA, and SA were significantly higher in the salt stress-sensitive genotype ZS11 than in the salt-resistant genotype Westar **(Fig. 10L-N)**, while in the roots, the concentrations of JA, SA, and BR were different between Westar and ZS11 **(Fig. 10O)**. Further, we also investigated the expression profiling of some genes involving the biosynthesis of stress-related phytohormones, such as ABA, JA, and SA, and found that most of them showed much higher transcriptional abundances in the shoots of Westar than in the shoots of ZS11 **(Supplementary Fig. S18)**. Based on all the results above, we proposed that ABA, JA, and SA might be involved in the pectin-mediated shoot salt resistance between Westar and ZS11.

**Fig. 10.**
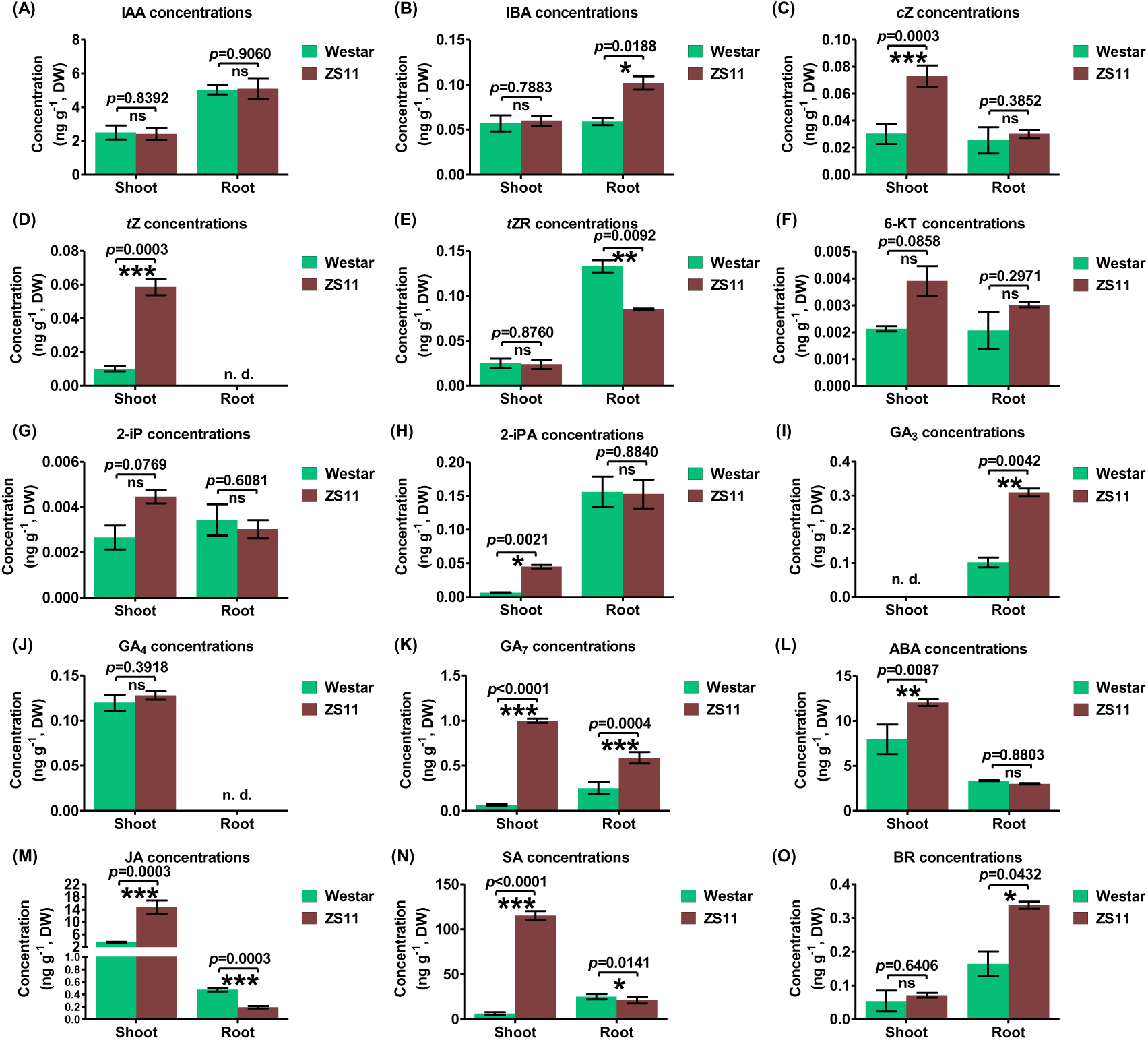
Comparative analysis of phytohormone profiling of Westar and ZS11 under salt stress. **(A-O)** Concentrations of IAA **(A)**, IBA **(B)**, *c*Z **(C)**, *t*Z **(D)**, *t*ZR **(E)**, 6-KT **(F)**, 2-iP **(G)**, 2-iPA **(H)**, GA3 **(I)**, GA4 **(J)**, GA7 **(K)**, ABA **(L)**, JA **(M)**, SA **(N)**, and BR **(O)** in the shoots and roots of Westar and ZS11 under salt stress. IAA, indole-3-acetic acid; IBA, indole-3-butyric acid; *c*Z, *cis*-Zeatin; *t*Z, *trans*-Zeatin; iP, isopentenyladenine; *t*ZR, trans-zeatin riboside; KT, kinetin; GA, gibberellic acid; ABA, abscisic acid; JA, jasmonic acid; SA, salicylic acid; BR, brassinosteroids. Uniform rapeseed plants after seed germination were grown hydroponically under NaCl-free conditions for 10 d, and then transferred to 200 mM NaCl for 5 d until sampling. Data are presented as the mean (n=5) ± s.d. Significant differences (*, *p* < 0.05; **, *p* < 0.01; ***, *p* < 0.001) were determined by unpaired two-tailed Student’s *t*-tests between two groups using the SPSS 17.0 toolkit. ns, not significant.

## Discussion

Salt stress resistance at the seedling stage is crucial for plant establishment and high yield in saline soils **(Wan et al., 2017)**. *B. napus* are relatively more resistant to salt stress as compared to their diploid relatives (e.g.) *B. campestris*, *B. nigra*, and *B. oleracea*; however, some studies have shown that even within amphitetraploids or diploids, genotypic variation exists with regard to salt stress resistance **(Shah et al., 2021)**. A previous study revealed that overexpression of two candidate genes predicted by genome-wide association studies in rapeseed, *BnCKX5* and *BnERF3*, were found to increase the sensitivity to salt stress at the germination stage **(Zhang et al., 2022)**. However, the regulatory mechanisms underlying these two genes regulating salt stress resistance and Na^+^ homeostasis remain to be elucidated in rapeseed. Dissecting the genetic basis of salt stress resistance provides valuable gene resources for improving the salt stress tolerance of *B. napus*.

### Molecular implications for the genetic improvement of salt stress resistance in the ZS11

ZS11 has been widely grown in China by virtue of its high seed yield production and good quality **(Sun et al., 2017)**. However, ZS11 is hypersensitive to salt stress **(Ibrahim et al., 2022; Wan et al., 2022)**, which severely hinders its growing in saline soils. Therefore, mining elite gene resources is very important for the genetic improvement of salt stress resistance in ZS11. Comparative omics analyses were conducted to reveal the molecular mechanism underlying differential salt resistance of between the salt-resistant cultivars Huayouza 62/Yangyou 9 and the salt-sensitive cultivar ZS11 **(Ibrahim et al., 2022; Wan et al., 2022)**. However, the core regulatory genes responsible for Na^+^homeostasis were not identified between the salt-resistant and salt-sensitive genotypes although some DEGs were predicted between these two genotypes.

Westar is widely used as a major transgenic receptor of rapeseed **(Cardoza and Stewart, 2003)**; however, it is hypersensitive to multiple abiotic and biotic stresses, such as such as boron deficiency **(Hua et al., 2016)** and blackleg disease **(Andreasson et al., 2001)**. Nevertheless, we found that Westar was a salt-resistant cultivar **(Fig. 1)**, which was also confirmed by a previous study **(Yong et al., 2015)**. Our previous studies showed that the differential resistance against boron deficiency and cadmium toxicity between Westar and ZS11 was related to their differences in the cell wall characteristics **(Hua et al., 2016; Zhang et al., 2019)**. In this study, we revealed that Westar outperformed than ZS11 under salt stress, and identified the core regulatory gene responsible for the differential salt stress resistance between the rapeseed genotypes through ionomics-assisted multiomics analysis. The dissection of the mechanisms underlying differential salt resistance between Westar and ZS11 not only provides valuable guidance for the selection of transgenic receptors when performing studies for salt resistance, but also is significant for the genetic improvement of salt stress resistance in ZS11.

### Pectin modification plays key roles in the resistance of Westar against salt stresses and other nutrient stresses

The plant cell wall is the first barrier between cell content and external salt. Plant cell walls consist of polysaccharides, various structural proteins, and fatty acid-derived compounds (such as cellulose, hemicelluloses, pectins, lignin, and suberin. The negative charge of cell walls is physiologically important for strengthening the cell wall via pectins cross-linking. However, under high salt concentrations, the surplus of Na^+^ could replace Ca^2+^, thus interrupting normal pectin cross-linking **(Dabravolski and Isayenkov, 2023)**.

In our previous study, we found that *BnaA8.PME3*, a *PME* homolog, might play a central role in the regulation of differential plant cadmium resistance between the cadmium-resistant genotype ZS11 and the cadmium-sensitive genotype Westar through modulating pectin-mediated cell wall cadmium retention **(Zhang et al., 2019)**. PMEs are novel and often overlooked players in the development of salt stress resistance. Previously, few PMEs have been functionally demonstrated to be involved in the regulation of salt stress resistance, particularly in the polyploid crop species. As yet, among the *PME* family genes, only *AtPME31* was found to act as a positive regulator of salt stress resistance, and knockdown of *PME31* resulted in the hypersensitivity of Arabidopsis plants to salt stress **(Yan et al., 2018)**. However, this study thought AtPME31 to be localized on the plasma membrane **(Yan et al., 2018)**, which might be attributed to the missing use of plasmolysis for the dissociation between the plasma membrane and the cell wall. In this study, through integrating fine isolation of cell wall components and characterization of ionomic, genomic, and transcriptomic profiling, we proposed pectin demethylation as to be a main factor responsible for differential salt stress resistance between Westar and ZS11. Further, we identified *BnaC9.PME47* as a positive regulator for salt stress resistance in rapeseed **(Figs. 7, 8)**, Moreover, *BnaC9.PME47* was identified to be localized in two reported QTL regions responsible for salt stress resistance in rapeseed **(Lang et al., 2017; Yong et al., 2015)**; moreover, it was strongly induced by salt stress, and presented nearly the highest expression abundance among the *BnaPME* family. All of the current results highlighted the potential pivotal importance of *BnaC9.PME47* in regulating salt stress resistance among the huge *PME* family in rapeseed. As yet, transcriptional regulation of *PMEs* remains largely unknown. GO and KEGG analysis of the DEGs between Westar and ZS11, identification of the *cis*-acting regulatory elements in the promoter region of *BnaC9.PME47*, and phytohomone profiling revealed that ABA and its responsive elements might be pivotal for the transcriptional regulation of *BnaC9.PME47*. Mutation of *atpme47* resulted in the hypersensitivity of Arabidopsis plants to salt stress **(Fig. 9)**, which indicated the key role of pectin demethylation-mediated cell wall Na^+^ retention in conferring salt stress resistance. Moreover, ionomic profiling showed that the low Na^+^ concentration in the Arabidopsis plants of *atpme47-2* still resulted in the loss of resistance against salt stress, which highlighted the essential role of subcellular Na^+^ distribution (including cell wall Na^+^ retention) in determining plant resistance under salt stress.

## Acknowledgement

We are very grateful to the editor and reviewers for critically evaluating the manuscript and providing constructive comments for its improvement.

## References

Almeida-Trapp M, Mithöfer A. Quantification of phytohormones by HPLC-MS/MS including phytoplasma-infected plants. Methods Mol Biol 2019, 1875:345–358.

Almeida Trapp M, De Souza GD, Rodrigues-Filho E, Boland W, Mithöfer A. Validated method for phytohormone quantification in plants. Front Plant Sci 2014, 5:417.

Andreasson E, Wretblad S, Granér G, Wu X, Zhang J, Dixelius C, Rask L, Meijer J. The myrosinase-glucosinolate system in the interaction between *Leptosphaeria maculans* and *Brassica napus*. Mol Plant Pathol 2001, 2:281–6.

Byrt CS, Munns R, Burton RA, Gilliham M, Wege S. Root cell wall solutions for crop plants in saline soils. Plant Sci 2018, 269:47–55.

Cardoza V, Stewart CN. Increased Agrobacterium-mediated transformation and rooting efficiencies in canola (*Brassica napus* L.) from hypocotyl segment explants. Plant Cell Rep 2003, 21:599–604.

Chen T, Liu D, Niu X, Wang J, Qian L, Han L, Liu N, Zhao J, Hong Y, Liu Y. Antiviral Resistance Protein Tm-2^2^ Functions on the Plasma Membrane. Plant Physiol 2017 Apr;173(4):2399–2410.

Dabravolski SA, Isayenkov SV. The regulation of plant cell wall organisation under salt stress. Front Plant Sci 2023, 14:1118313.

Deng S, Sun J, Zhao R, et al. 2015. *Populus euphratica* APYRASE2 enhances cold tolerance by modulating vesicular trafficking and extracellular ATP in Arabidopsis plants. Plant Physiol 169, 530–548.

Dittmer J, Lusseau T, Foissac X, Faoro F. 2021. Skipping the insect vector: plant stolon transmission of the phytopathogen *’Ca Phlomobacter fragariae’* from the Arsenophonus clade of insect endosymbionts. Insects 12, 93.

Fang C, Li K, Wu Y, Wang D, Zhou J, Liu X, Li Y, Jin C, Liu X, Mur LAJ, Luo J. OsTSD2-mediated cell wall modification affects ion homeostasis and salt tolerance. Plant Cell Environ 2019, 42:1503–1512.

Feng W, Kita D, Peaucelle A, Cartwright HN, Doan V, Duan Q, Liu MC, Maman J, Steinhorst L, Schmitz-Thom I, Yvon R, Kudla J, Wu HM, Cheung AY, Dinneny JR. The FERONIA receptor kinase maintains cell-wall integrity during salt stress through Ca^2+^ signaling. Curr Biol 2018, 28:666–675.

Feng ZT, Deng YQ, Zhang SC, et al. K (^+^) accumulation in the cytoplasm and nucleus of the salt gland cells of Limonium bicolor accompanies increased rates of salt secretion under NaCl treatment using NanoSIMS. Plant Sci 2015, 238:286–296.

Higo K, Ugawa Y, Iwamoto M, Korenaga T. 1999. Plant *cis*-acting regulatory DNA elements (PLACE) database: 1999. Nucleic Acids Research 27, 297–300.

Hocq L, Pelloux J, Lefebvre V (2017) Connecting homogalacturonan-type pectin remodeling to acid growth. Trends Plant Sci 22: 20–29.

Hua YP, Zhang D, Zhou T, He M, Ding G, Shi L, Xu F. Transcriptomics-assisted quantitative trait locus fine mapping for the rapid identification of a nodulin 26-like intrinsic protein gene regulating boron efficiency in allotetraploid rapeseed. Plant Cell Environ 2016, 39:1601–18.

Hua YP, Zhang YF, Zhang TY, Chen JF, Song HL, Wu PJ, Yue CP, Huang JY, Feng YN, Zhou T. Low iron ameliorates the salinity-induced growth cessation of seminal roots in wheat seedlings. Plant Cell Environ 2023, 46:567–591.

İbrahimova U, Kumari P, Yadav S, Rastogi A, Antala M, Suleymanova Z, Zivcak M, Tahjib-Ul-Arif M, Hussain S, Abdelhamid M, Hajihashemi S, Yang X, Brestic M. Progress in understanding salt stress response in plants using biotechnological tools. J Biotechnol 2021, 329:180–191.

Kanehisa M, Sato Y. 2020. KEGG Mapper for inferring cellular functions from protein sequences. Protein Science 29, 28–35.

Kanneganti V, Gupta AK. Isolation and expression analysis of OsPME1, encoding for a putative Pectin Methyl Esterase from *Oryza sativa* (subsp. *indica*). Physiol Mol Biol Plants 2009, 15:123–31.

Kaur N, Sharma I, Kirat K. and Pati PK. Detection of reactive oxygen species in *Oryza sativa* L. (Rice). Bio-protocol 2016, 6:e2061.

Kotula L, Garcia Caparros P, Zörb C, Colmer TD, Flowers TJ. 2020. Improving crop salt tolerance using transgenic approaches: An update and physiological analysis. Plant, Cell and Environment 43, 2932–2956.

Lang L, Xu A, Ding J, Zhang Y, Zhao N, Tian Z, Liu Y, Wang Y, Liu X, Liang F, Zhang B, Qin M, Dalelhan J, Huang Z. Quantitative trait locus mapping of salt tolerance and identification of salt-tolerant genes in *Brassica napus* L. Front Plant Sci 2017, 8:1000.

Lescot M, Déhais P, ThijsG, Marchal K, MoreauY, Van de PeerY, Rouzé P, Rombauts S. 2002. PlantCARE, a database of plant *cis*-acting regulatory elements and a portal to tools for *in silico* analysis of promoter sequences. Nucleic Acids Research 30, 325–327.

Liu Y, Yu Y, Sun J, Cao Q, Tang Z, Liu M, Xu T, Ma D, Li Z, Sun J. Root-zone-specific sensitivity of K^+^-and Ca^2+^-permeable channels to H_2_O_2_ determines ion homeostasis in salinized diploid and hexaploid *Ipomoea trifida*. J Exp Bot 2019, 70:1389–1405.

Livak KJ, Schmittgen TD. 2001. Analysis of relative gene expression data using real-time quantitative PCR and the 2^-ΔΔC^*_T_* method. Methods 25, 402–408.

Louvet R, Cavel E, Gutierrez L, Guénin S, Roger D, Gillet F, Guerineau F, Pelloux J. Comprehensive expression profiling of the pectin methylesterase gene family during silique development in Arabidopsis thaliana. Planta 2006, 224:782–91.

Lv S, Jiang P, Chen X, Fan P, Wang X, Li Y. 2012. Multiple compartmentalization of sodium conferred salt tolerance in *Salicornia europaea*. Plant Physiology and Biochemistry. 51, 47–52.

Maillard A, Etienne P, Diquélou S, Trouverie J, Billard V, Yvin JC, Ourry A. 2016. Nutrient deficiencies in *Brassica napus* modify the ionomic composition of plant tissues: a focus on cross-talk between molybdenum and other nutrients. Journal of Experimental Botany 67, 5631–5641.

Mi H, Muruganujan A, Thomas PD. 2013. PANTHER in 2013: modeling the evolution of gene function, and other gene attributes, in the context of phylogenetic trees. Nucleic Acids Research 41, D377–D386.

Mohamed I, Shalby N, Mahmoud E, Batool M, Wang C, Wang Z, Salah A, Rady M, Jie K, Wang B, Zhou GS. RNA-seq analysis revealed key genes associated with salt tolerance in rapeseed germination through carbohydrate metabolism, hormone, and MAPK signaling pathways. Industrial Crops and Products 2022, 176:114262.

Ouyang SQ, Liu YF, Liu P, et al. Receptor-like kinase OsSIK1 improves drought and salt stress tolerance in rice (*Oryza sativa*) plants. Plant J 2010, 62:316–329.

Shah AN, Tanveer M, Abbas A, Fahad S, Baloch MS, Ahmad MI, Saud S, Song Y. Targeting salt stress coping mechanisms for stress tolerance in Brassica: A research perspective. Plant Physiol Biochem 2021, 158:53–64.

Song JM, Guan Z, Hu J, Guo C, Yang Z, Wang S, Liu D, Wang B, Lu S, Zhou R, Xie WZ, Cheng Y, Zhang Y, Liu K, Yang QY, Chen LL, Guo L. Eight high-quality genomes reveal pan-genome architecture and ecotype differentiation of *Brassica napus*. Nat Plants. 2020, 6:34–45.

Sun F, Fan G, Hu Q, Zhou Y, Guan M, Tong C, Li J, Du D, Qi C, Jiang L, Liu W, Huang S, Chen W, Yu J, Mei D, Meng J, Zeng P, Shi J, Liu K, Wang X, Wang X, Long Y, Liang X, Hu Z, Huang G, Dong C, Zhang H, Li J, Zhang Y, Li L, Shi C, Wang J, Lee SM, Guan C, Xu X, Liu S, Liu X, Chalhoub B, Hua W, Wang H. The high-quality genome of *Brassica napus* cultivar ’ZS11’ reveals the introgression history in semi-winter morphotype. Plant J. 2017, 92(3):452–468.

Wan H, Chen L, Guo J, Li Q, Wen J, Yi B, Ma C, Tu J, Fu T, Shen J. Genome-wide association study reveals the genetic architecture underlying salt tolerance-related traits in rapeseed (*Brassica napus* L.). Front Plant Sci 2017, 8:593.

Wan H, Qian J, Zhang H, Lu H, Li O, Li R, Yu Y, Wen J, Zhao L, Yi B, Fu T, Shen J. Combined transcriptomics and metabolomics analysis reveals the molecular mechanism of salt tolerance of Huayouza 62, an elite cultivar in rapeseed (*Brassica napus* L.). Int J Mol Sci 2022, 23:1279.

Woodward AW, Bartel B. Auxin: regulation, action, and interaction. Ann Bot. 2005, 95:707–35.

Yan J, He H, Fang L, Zhang A. Pectin methylesterase 31 positively regulates salt stress tolerance in Arabidopsis. Biochem Biophys Res Commun 2018, 496:497–501.

Yang HL, Liu J, Huang SM, Guo TT, Deng LB, Hua W. 2014. Selection and evaluation of novel reference genes for quantitative reverse transcription PCR (qRT-PCR) based on genome and transcriptome data in *Brassica napus* L. Gene 538, 113–122.

Yang Z, Nielsen R. 2000. Estimating synonymous and nonsynonymous substitution rates under realistic evolutionary models. Molecular Biology and Evolution 17, 32–43.

Yang Z, Wang C, Xue Y, et al. Calcium-activated 14-3-3 proteins as a molecular switch in salt stress tolerance. Nat Commun 2019, 10:1199.

Yong HY, Wang C, Bancroft I, Li F, Wu X, Kitashiba H, Nishio T. Identification of a gene controlling variation in the salt tolerance of rapeseed (*Brassica napus* L.). Planta 2015, 242:313–26.

Zhang G, Zhou J, Peng Y, Tan Z, Li L, Yu L, Jin C, Fang S, Lu S, Guo L, Yao X. Genome-wide association studies of salt tolerance at seed germination and seedling stages in *Brassica napus*. Front Plant Sci 2022, 12:772708.

Zhang ZH, Zhou T, Tang TJ, Song HX, Guan CY, Huang JY, Hua YP. A multiomics approach reveals the pivotal role of subcellular reallocation in determining rapeseed resistance to cadmium toxicity. J Exp Bot 2019, 70:5437–5455.

Zhao KF, Fan H, Zhou S, Song J. 2003. Study on the salt and drought tolerance of *Suaeda salsa* and *Kalanchoe claigremontiana* under iso-osmotic salt and water stress. Plant Science 165, 837–844.

Zhou T, Hua Y, Zhang B, Zhang X, Zhou Y, Shi L, Xu F. Low-boron tolerance strategies involving pectin-mediated cell wall mechanical properties in *Brassica napus*. Plant Cell Physiol 2017, 58:1991–2005.

Zhou T, Yue CP, Huang JY, Cui JQ, Liu Y, Wang WM, Tian C, Hua YP. 2020. Genome-wide identification of the amino acid permease genes and molecular characterization of their transcriptional responses to various nutrient stresses in allotetraploid rapeseed. BMC Plant Biology 20, 151.

Zhou T, Yue CP, Liu Y, Zhang TY, Huang JY, Hua YP. Multiomics reveal pivotal roles of sodium translocation and compartmentation in regulating salinity resistance in allotetraploid rapeseed. J Exp Bot 2021, 72:5687–5708.

Zhu XF, Wang ZW, Wan JX, Sun Y, Wu YR, Li GX, Shen RF, Zheng SJ. Pectin enhances rice (*Oryza sativa*) root phosphorus remobilization. J Exp Bot 2015, 66:1017–24.

